# Reducing module size bias of participation coefficient

**DOI:** 10.1101/747162

**Authors:** Mangor Pedersen, Amir Omidvarnia, James M. Shine, Graeme D. Jackson, Andrew Zalesky

## Abstract

Both natural and engineered networks are often modular. Whether a network node interacts with only nodes from its own module or nodes from multiple modules provides insight into its functional role. The participation coeffcient (*PC*) is typically used to measure this attribute although its value also depends on the size of the module it belongs to, often leading to non-intuitive identification of highly connected nodes. Here, we develop a normalized *PC* that overcomes the module size bias associated with the conventional *PC*. Using brain, *C.elegans*, airport and simulated networks, we show that our measure of participation alleviates the module size bias, while preserving conceptual and mathematical properties, of the classic formulation of *PC*. Unlike the conventional *PC*, we identify London and New York as high participators in the air traffic network and demonstrate stronger associations with working memory in human brain networks, yielding new insights into nodal participation across network modules.

## Introduction

Many natural and engineered networks are modular [1–3]. Networks that are highly modular can be partitioned into communities of nodes, or modules, such that the density of connections is greater between the nodes within modules, relative to the density between nodes in different modules. Some nodes have connections that are distributed across many modules, whereas others are only connected with other nodes in their own module. This distinction can provide important insight into a node’s functional role in a modular architecture.

A node’s inter-modular connectivity is typically quantified with the participation coeffcient (*PC*) [4]. *PC* provides insights into how specific nodes communicate between modules in a range of real-world networks, including air traffic and brain networks [5–13]. To compute a node’s *PC*, the proportion of a node’s connections to each module is first determined, yielding a proportion for each module. These proportions are then squared, summed across all modules and the resulting summand is subtracted from one to yield the node’s *PC*. *PC* of zero indicates a node that only connects with other nodes in its own module, whereas nodes with connections that are uniformly distributed across all modules have *PC* that approaches one.

*PC* tacitly assumes that all modules in a network are equally sized. This is, however, rarely the case in real-world networks. *PC* is consequently susceptible to a bias wherein nodes in small modules often have high *PC* and nodes in large modules often have low *PC* [14]. This bias is exemplified in the two networks shown in Fig. 1, where node *i* in network B, by the virtue of belonging to a larger module, has a lower *PC* than node *i* in network A, even though node *i* has the same inter-modular connectivity in both networks. We refer to this as the module size bias of *PC*, which we show can yield biased inference about a node’s participation.

**Figure 1:**
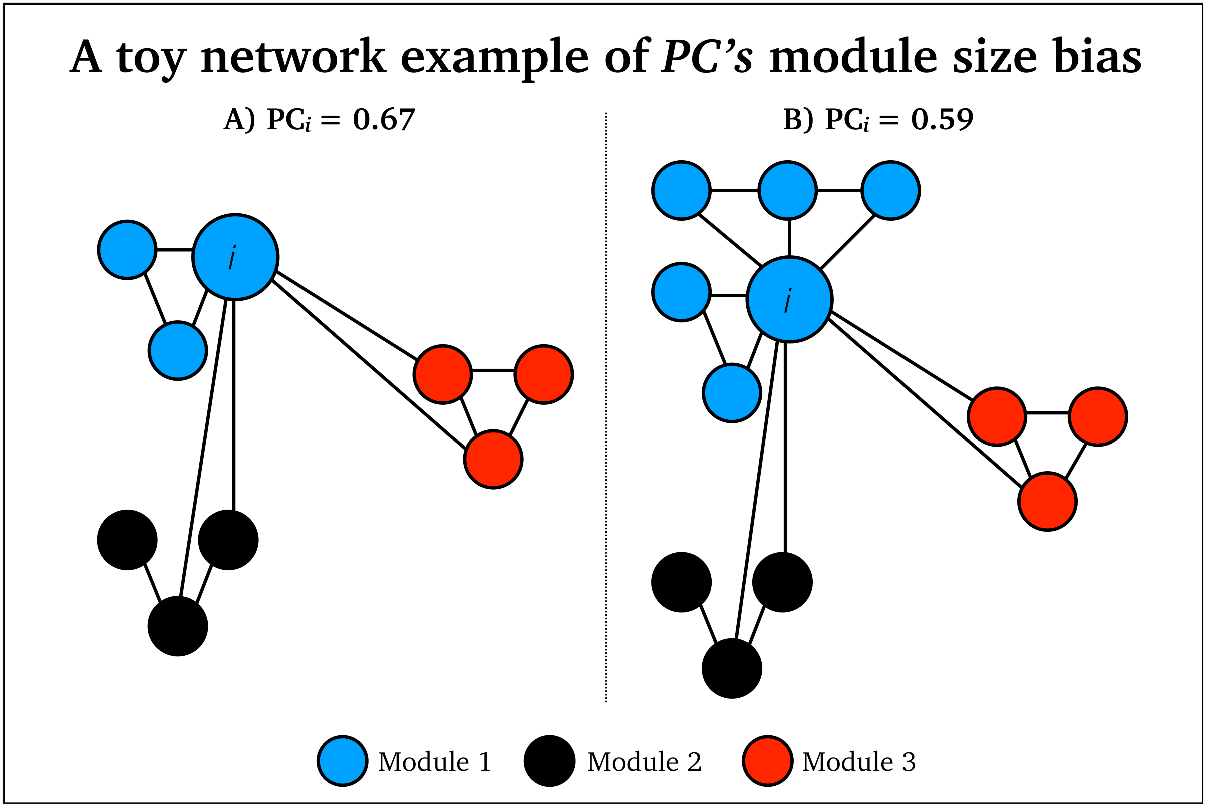
Toy network examples of *PC’s* module size bias: Node *i* has identical intermodule connectivity in network A and B, but it has lower *PC*_*i*_ in network B as it belongs to a larger blue module than network A.

The module size bias of *PC* was first highlighted by Klimm and others [14]. These authors proposed a dispersion index as an alternative to *PC*. In this paper, we propose a modification of the original *PC* measure rather than constructing a new measure of network integration. Our aim is to reduce the module size bias of *PC* while retaining the underlying mathematical assumptions, and numerical range, of the original *PC* measure. To achieve this, we developed a normalized *PC* in which a node’s participation is benchmarked to an ensemble of random networks matched in node degree and connection density [15]. Our normalized participation coeffcient (*PC*_*norm*_) accounts for the (intra-modular) connectivity expected by chance, given the size of network modules. To validate *PC*_*norm*_, we used three real-world networks: i) undirected functional MRI (fMRI) brain network data with 100 brain regions from 1003 healthy adults (ages 22-35) participating in the Human Connectome Project [16]; ii) a single directed neural network from the *C.elegans* nematode containing a total of 277 neurons [17,18]; and iii) a single directed flight network with 500 airports [19], as well as simulated networks [4].

Firstly, simulations were undertaken to test the hypothesis that *PC*_*norm*_ preserves the network features and numerical range of *PC*, in networks where all modules were equally sized. We then hypothesized that *PC*_*norm*_ would reduce the module size bias inherent to *PC* in real-world networks across a range of spatial resolution parameters and network density thresholds. In addition to our main hypotheses, we assessed nodal traits of *PC*_*norm*_, to evaluate their relevance in neurobiological and air-traffic systems. We also investigated whether *PC*_*norm*_ measured in fMRI networks associated with behavior more strongly than *PC*. Together, our results demonstrate a simple refinement of *PC* that retains the interpretation of *PC* in real-world networks but alleviates the module size bias. Unlike the conventional *PC*, we identify London (LHR) and New York (JFK), among other airports, as high participators in the air traffic network, and demonstrate stronger associations with working memory in human brain networks. We hope *PC*_*norm*_ will lead to a more reliable estimation of nodal integration in complex networks.

## Results

### Normalized participation coeffcient (*PC*_*norm*_)

The participation coeffcient (*PC*) measures whether a node interacts with only nodes from its own module or nodes from multiple modules [4]. Formally, *PC* of node *i* is given by,

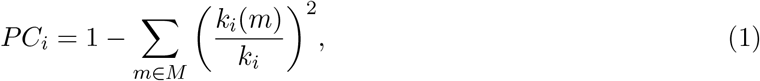

where *M* is the set of network modules, *k*_*i*_(*m*) is the degree between node *i* and all nodes in module *m* and *k_i_* is the degree between node *i* and all other nodes in the entire network. A node with *PC* of zero only interacts with nodes comprising its own module, while nodes with connections uniformly distributed across all modules have a *PC* that approaches one. As demonstrated below and discussed elsewhere [14], *PC* is limited by an inherent module size bias, which may lead to inaccurate inference in networks with modules that vary in size.

To alleviate the module size bias, we propose the normalized participation coeffcient (*PC*_*norm*_),

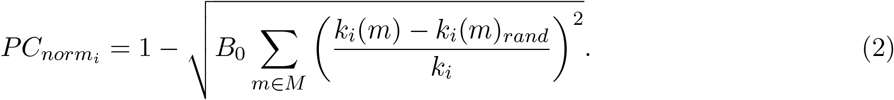

The key difference between *PC* and *PC*_*norm*_ of node *i* is subtraction of the normalization factor, *k*_*i*_(*m*)_*rand*_, from *k*_*i*_(*m*). This normalization factor denotes the median intra-modular degree for node *i* across randomized networks generated with an established network rewiring algorithm that preserves connection density and node degree (i.e., *k*_*i*_). This also means that *k*_*i*_ is the same for original and randomized networks [15]. We found that 1000 network randomizations were adequate to return a stable estimate of *PC*_*norm*_ (see Supplementary Fig 1). In each randomized network, all edges were rewired five times. We used the same underlying modular network structure (*M*) for original *k*_*i*_(*m*) and random *k*_*i*_(*m*)_*rand*_. To constrain the range of *PC*_*norm*_ between 0 (low network integration) and 1 (high network integration), we add the multiplicative term *B*_0_ = 0.5, and we also calculate the square root of the difference of participation between original and randomized networks.

### *PC* and *PC*_*norm*_ are comparable in simulated networks without a module size bias

Firstly, we aimed to establish that our normalization process does not alter the conceptual basis of the *PC*, or its mathematical properties and relationships with other topological attributes. To discount this possibility, we implemented the network simulation described by Guimerà and Amaral [4] and verified that *PC* and *PC*_*norm*_ yielded comparable values in a network comprising equally-sized modules. Specifically, we simulated binary networks with 100 nodes, and four equal-sized modules (each containing 25 nodes). Given that the modules were equally sized, module size bias was absent by design, and thus *PC* and *PC*_*norm*_ should not markedly differ. We added connections either within or between the different modules through a range of probabilities (0 to 1, in increments of 0.01), and calculated *PC*_*norm*_ (Fig. 2A), *PC* (Fig. 2B), *PC*_*norm*_ minus *PC* (Fig. 2C) and modularity Q-score (Fig. 2D), for all possible connectivity probability values.

**Figure 2:**
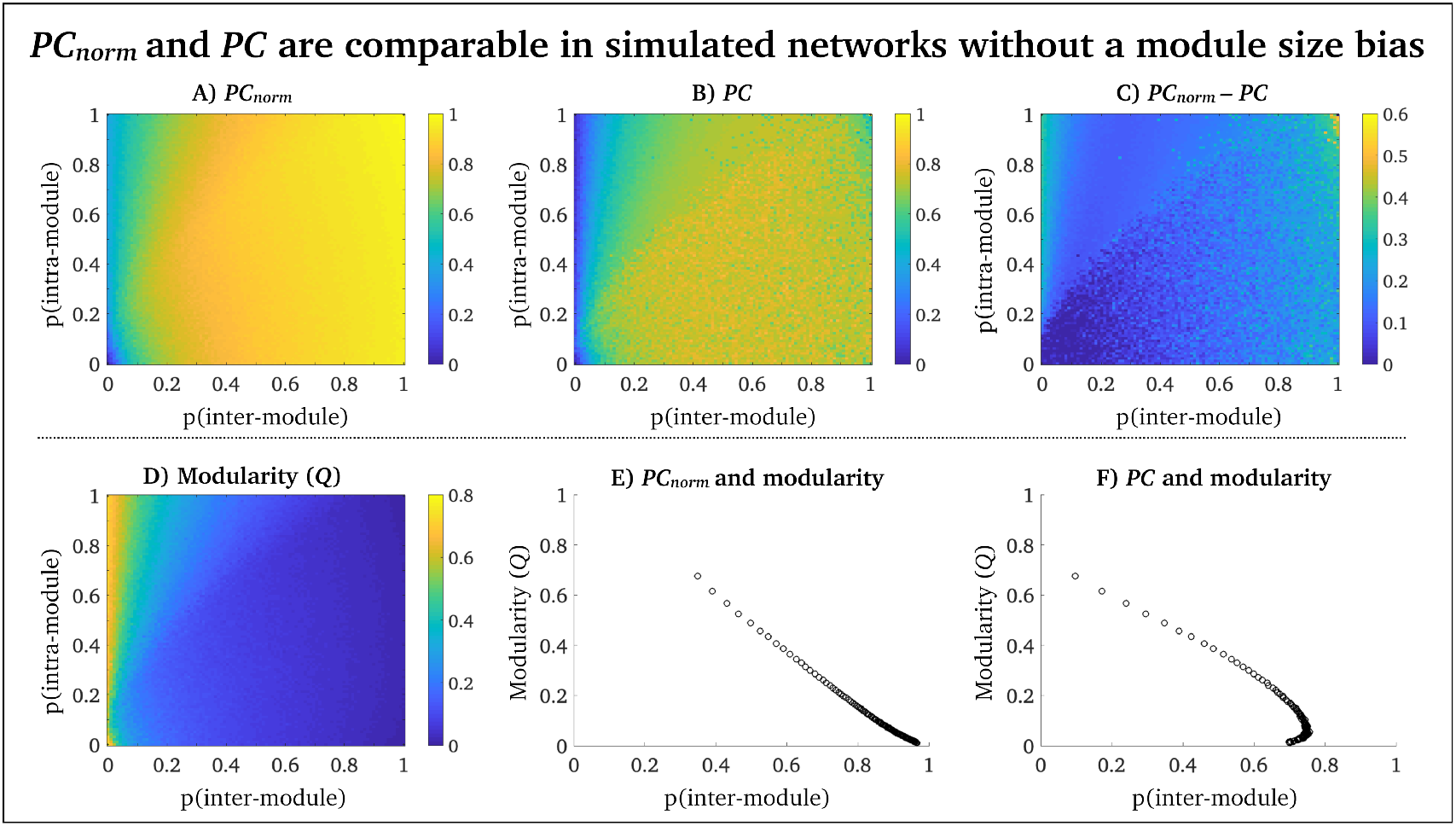
Simulations in networks without module size bias: Simulation results for *PC*_*norm*_ and *PC* in networks with equally sized modules, where the heat maps show that average *PC* across all nodes. (A) *PC*_*norm*_ across probabilities of intra- (*y*-axis) and inter-module (*x*-axis) connectivity. (B) *PC* across probabilities of intra- (*y*-axis) and inter-module (*x*-axis) connectivity. (C) *PC*_*norm*_ minus *PC* across probabilities of intra- (*y*-axis) and inter-module (*x*-axis) connectivity. (D) modularity (Q-score) across probabilities of intra- (*y*-axis) and inter-module (*x*-axis) connectivity. (E) average modularity for *PC*_*norm*_ across inter-module (*x*-axis) connectivity. (F) average modularity for *PC* across inter-module (*x*-axis) connectivity. This simulation suggests that *PC*_*norm*_ preserves the mathematical, and conceptual, features of the original *PC* measure.

Figure 2A and B suggest that *PC*_*norm*_ and *PC* yield highly comparable estimates across a range of simulated networks with equally sized modules. More specifically, the relationship between *PC*_*norm*_ and *PC* was strong across all simulated networks (Pearson’s *r* = 0.91). This suggests that *PC*_*norm*_ remains consistent with *PC* in networks with no module size bias, despite our algorithmic changes to node participation. Our simulation additionally shows that *PC*_*norm*_ values reside within the same range as *PC* (i.e., [0,1]), although *PC*_*norm*_ tends to have higher values than *PC* (see Fig 2, 3 and 8). *PC*_*norm*_ is also shows a more linearly inverse relationship between modularity and inter-module connectivity than *PC* (Fig. 2E-F). This phenomena was also outlined by Guimerà and Amaral in their seminal *PC* paper [4], where the original PC algorithm may be unstable if a node has more than 50% inter-modular connections. This suggests that *PC*_*norm*_ may be an appropriate metric in networks with numerous inter-modular connections.

**Figure 3:**
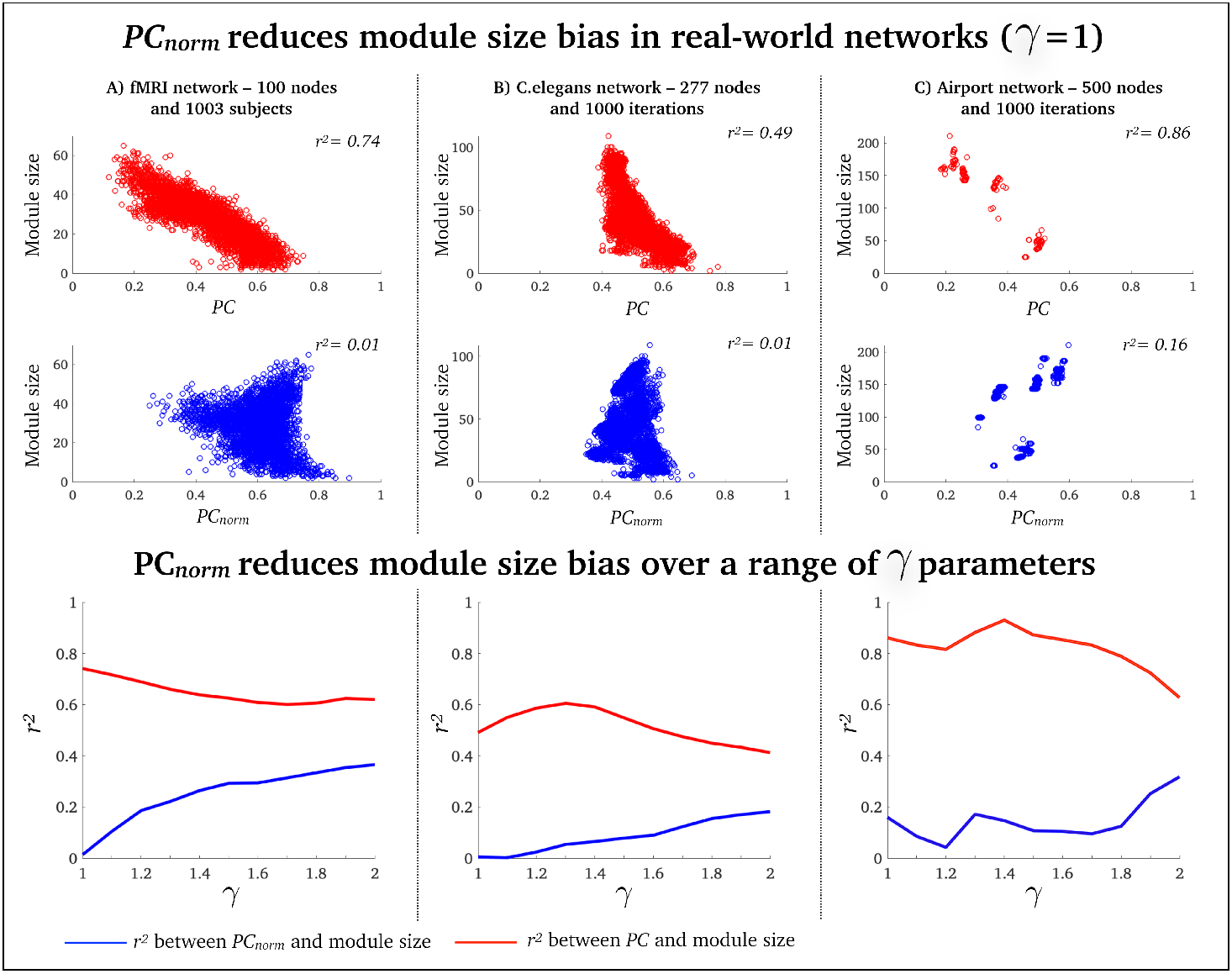
*PC*_*norm*_ reduces *PC’s* module size bias: Relationship between *PC* and module size (top row), *PC*_*norm*_ and module size (middle row), for *γ* = 1 (few and large module). Each data point is the average participation of all nodes within a module. Displayed are (A) 3357 modules (data points) across 1003 fMRI networks; (B) 6020 modules (data points) across 1000 *C.elegans* networks and (C); 3998 modules (data points) across 1000 airport networks. The scatter plots for the airport network appear sparser due to greater consistency across runs of the modular decomposition algorithm, relative to the other networks (see Supplementary Fig. 2). The bottom row shows the mean difference between *PC*, *PC*_*norm*_ and module size for *γ* between 1 and 2 in 0.1 increments. It is worth noting that *PC*’s module size bias is reduced for high *γ* values (*γ* > 1.5), where networks have numerous small modules with less variation in size, although *γ* > 1.5 is rarely used in complex network analysis [20].

### *PC*_*norm*_ reduces *PC’s* module size bias in real-world modular networks

Next, we aimed to test whether *PC*_*norm*_ alleviates the module size bias of *PC* and thus yields more intuitive conclusions about the importance of single nodes in three real-world networks. Using a Louvain modularity decomposition method with a spatial resolutions of *γ* = 1 [21] (*γ* = 1 yields few and large modules), we found that fMRI human brain networks had an average modularity Q-score of 0.35 ± 0.03 [SD]^1^, and an average of 3.52 ± 0.63 [SD] modules, across 1003 healthy adults; the *C.elegans* network had an average modularity Q-score of 0.42 ± 0.01 [SD], and an average of 6.02 ± 0.55 [SD] modules, across 1000 Louvain modularity iterations; the airport network had an average modularity Q-score of 0.50 ± 0.01 [SD] and an average of 4.01 ± 0.05 [SD] modules, across 1000 Louvain modularity iterations (see Supplementary Fig. 3, for results across a range of *γ* parameters). This resulted in a total of 3003 networks for analysis, where we for each network calculated the average *PC*_*norm*_ and *PC* across all (non-zero) nodes in each module. In our main analysis we included 11 values for the *γ*-parameters (1 to 2, in increments of 0.1). This was done to verify that *PC*_*norm*_ reduces *PC*’s module size bias across several resolution parameters and module sizes.

To enable statistical inference between the two squared correlation coefficients of interest (correlation value #1 = *r*^2^ between *PC* and module size; correlation value #2 = *r*^2^ between *PC*_*norm*_ and module size), we employed a bootstrapping approach using 10000 bootstrap samples with replacement [22]. Here, we aimed to test the null hypothesis of equality in the two correlation coefficients. For each sample, we computed squared correlation coefficients based on the pooled bootstrap samples of participation (*PC*_*norm*_ and *PC*) and module size, as well as its 95% confidence intervals (95% CI).

In line with our hypothesis, we found that the variance explained (*r*^2^) between the participation coefficient and module size was substantially lower for *PC*_*norm*_ relative to *PC*, where a module’s size was determined by the number of nodes it comprised. This indicates that *PC*_*norm*_ alleviates the module size bias of *PC*. We display correlation patterns between *PC*_*norm*_, *PC* and module size, in Fig. 3, and we summarize the statistical differences between *PC*_*norm*_, *PC* and module size in Table 1.

**Table 1:**
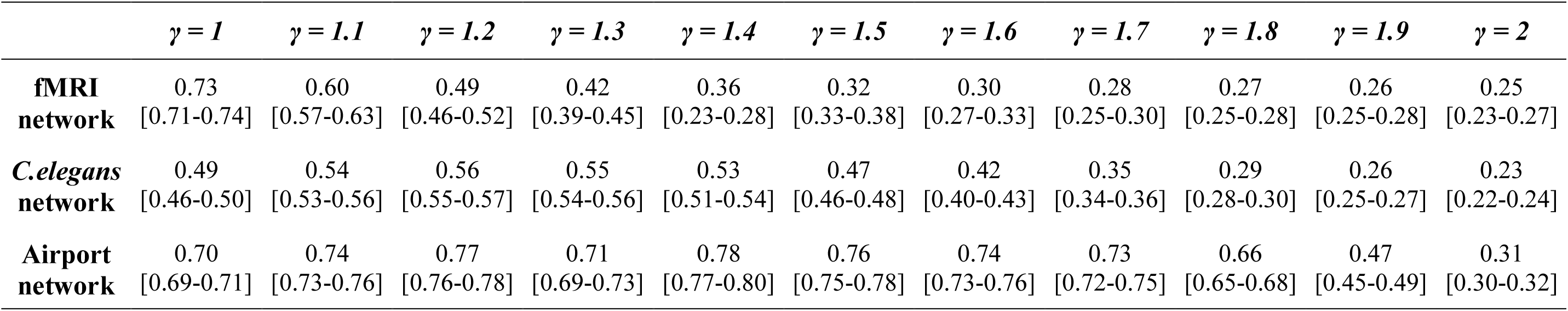
Mean r^2^ bootstrap difference, and 95% CI, between PC and PC_norm_ (correlation with module size)

Given that connection density – i.e., the proportion of edges in a network– can markedly impact a network’s modular architecture as well as other topological properties [23], we also confirmed that *PC*_*norm*_ reduces *PC*’s module size bias in fMRI networks, across multiple connection densities (Supplementary Fig. 4). From now on, we report results associated with *γ* = 1.

### Node-wise features of *PC*_*norm*_ in real-world modular networks

Next, we investigated the distinct inferences and conclusions that can be reached about the role of specific nodes when alleviating the module size bias. For this purpose, we subtracted average *PC* from *PC*_*norm*_ across 1003 fMRI network subjects, and 1000 modularity runs for *C.elegans* and airport networks. In the next few paragraphs we have summarized node-wise results for fMRI, *C.elegans* and airport networks. We have also provided a full set of node-wise results for *PC*_*norm*_ and *PC* in Supplementary Table 1, 2 and 3, including *z*-score differences between these two measures. A *z*-score was used to calculate the relative differences between *PC*_*norm*_ and *PC* because *PC*_*norm*_ tends to yield higher values than *PC* (Fig. 2 and 3). In Supplementary Fig. 5, we display the average modularity structure for fMRI, *C.elegans* and airport networks.

In the brain, the largest difference between *PC*_*norm*_ and *PC* was observed in cerebellum subregions crus I (z = 1.76), crus II (*z*-score = 1.56) and lobule VI (*z*-score = 1.42), but also in cortical areas such as inferior parietal cortex (*z*-score = 1.31) and superior frontal gyrus (z = 1.12) (Fig. 4A), areas that are known to connect to a variety of sub-networks in the brain [24], and that subserve higher order cognitive function [11]. Brain regions with the highest *PC*_*norm*_ were cerebellum lobule VI (*PC*_*norm*_ = 0.73), cerebellum crus I (*PC*_*norm*_ = 0.72; *PC* = 0.49), inferior parietal cortex (*PC*_*norm*_ = 0.71; *PC* = 0.46), middle frontal gyrus (*PC*_*norm*_ = 0.71; *PC* = 0.50) and precuneus (*PC*_*norm*_ = 0.70; *PC* = 0.52) (Supplementary Table 1).

**Figure 4:**
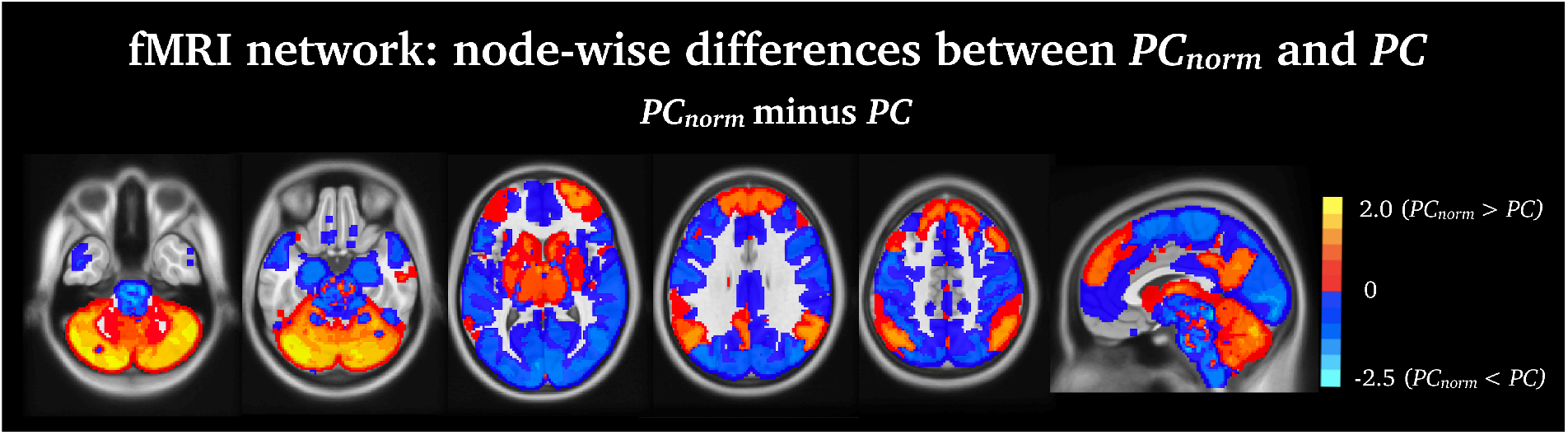
Node-wise fMRI differences between *PC*_*norm*_ and *PC:* Displayed are the subtracted group-average *PC* from *PC*_*norm*_ in fMRI networks (z-score). The largest difference between *PC*_*norm*_ and *PC* was observed in cerebellum subregions lobule VI, crus I and II, fronto-parietal and insular cortex.

For the *C.elegans* network, the greatest nodal differences between *PC*_*norm*_ and *PC* was observed in the largest module of the nematode encompassing inhibitory GABAergic neurons involved in head movement (RMDDL – *z*-score = 2.48; RICR – *z*-score = 2.35; RMDVL – *z*-score = 2.27; RMDDR – *z*-score = 2.26; and RMDVR *z*-score = 2.23 – Fig 5 and Supplementary Table 2). These were not the same subset of neurons with highest node participation. The strongest *PC*_*norm*_ was observed in several interneurons, also located in the head of the nematode including AVER (*PC*_*norm*_ = 0.87; *PC* = 0.75); AVAR (*PC*_*norm*_ = 0.85; *PC* = 0.75); AVAL (*PC*_*norm*_ = 0.84; *PC* = 0.75); AVBR (*PC*_*norm*_ = 0.83; *PC* = 0.74) and AVBL (*PC*_*norm*_ = 0.80; *PC* = 0.74), responsible for locomotor behavior, which is an important function for the *C.elegans* nematode. These locomotor interneurons have been found to be densely inter-connected across a range of sub-networks forming a selective ‘rich-club’ responsible for a bulk of neural signalling in the nervous system of the *C.elegans* nematode [13, 25–27]. See https://www.wormatlas.org/neurons/Individual%20Neurons/Neuronframeset.html, for full naming, definition and function of specific *C.elegans* neurons.

**Figure 5:**
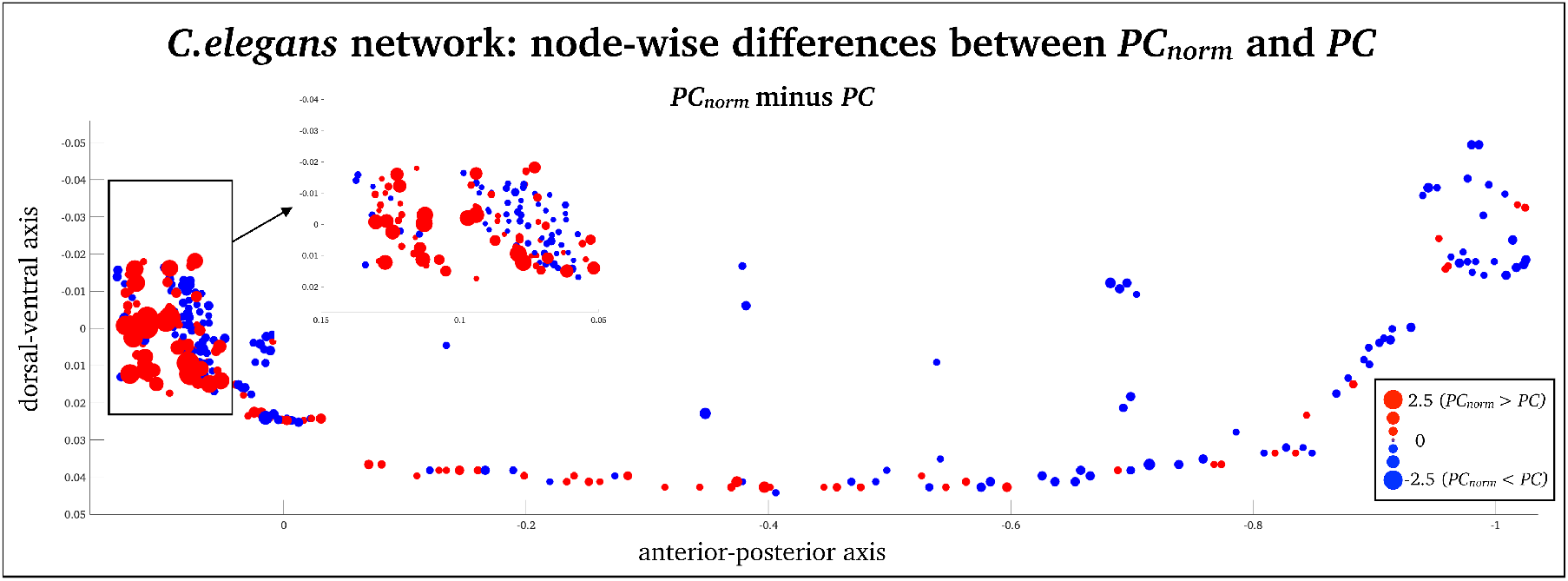
Node-wise *C.elegans* differences between *PC*_*norm*_ and *PC:* Z-score difference beteween *PC* and *PC*_*norm*_, in the *C.elegans* network (z-score). The largest difference between *PC*_*norm*_ and *PC* was observed in in inhibitory GABAergic neurons that belong to the largest module of the nematode encompassing RMDDL, RICR, RMDVL, RMDDR and RMDVR neurons.

In the airport network, the greatest difference between *PC*_*norm*_ and *PC* was observed in South American airports (Montevideo – *z*-score = −2.26; Sao Paolo – *z*-score = −2.22; Lima – *z*-score = −2.20; Rio de Janeiro – *z*-score = −2.19; and Buenos Aires – *z*-score = −2.18 – see Fig. 6A and Supplementary Table 3), which was the smallest module in this network. Notably, 8 of the 10 airports with highest *PC* belonged to the module encompassing Central and South America (Fig 6B – red circles). This was in contrast to the nodes with strongest *PC*_*norm*_, which included major airport hubs, including New York-JFK (*PC*_*norm*_ = 0.85; *PC* = 0.61), Punta Cana (*PC*_*norm*_ = 0.83; *PC* = 0.58), Toronto (*PC*_*norm*_ = 0.81; *PC* = 0.55), Montreal (*PC*_*norm*_ = 0.81; *PC* = 0.49) and Frankfurt (*PC*_*norm*_ = 0.78; *PC* = 0.62) (see blue circles in Fig. 6). These airports are known to have many intercontinental flights. This node-wise analysis suggests that *PC* overestimates the integrative nature of nodes within relatively small modules and that *PC*_*norm*_ can alleviate systematic module size bias, thereby enabling clearer and more intuitive conclusions to be drawn about the role of nodes.

**Figure 6:**
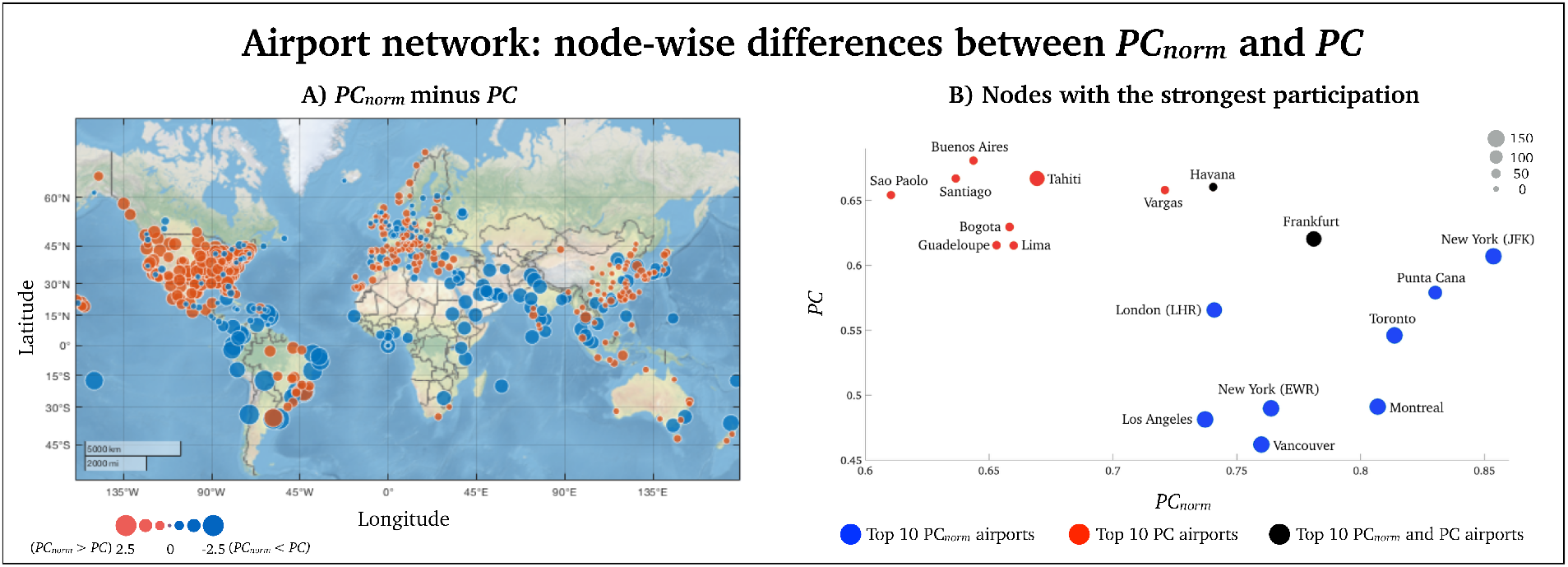
Node-wise difference between *PC*_*norm*_ and *PC* in the network of airports: (A) A world map displaying all airports included in the study. As highlighted in the manuscript, we observed the greatest difference between the two metrics in Central- and South American airports (red moudle). (B) A visual representation of the ten nodes with strongest *PC*_*norm*_ on the *x*-axis (blue nodes) and *PC* (red nodes) on the *y*-axis. The two black nodes were high for *PC*_*norm*_ and *PC*. Here, the size of each node is proportional to its module size (number of nodes in modules).

### *PC*_*norm*_ is more strongly correlated with working-memory performance than *PC*

Having established that *PC*_*norm*_ alleviates the module size bias of *PC* and yields a more parsimonious interpretation of node roles, we next aimed to test whether inter-individual variation in *PC*_*norm*_ measured in brain networks would associate more strongly with behavioral measures than *PC* [28].

We conducted an analysis informed by our previous study where we showed that inter-modular network switching was related to the following behavioral domains [8]: i) *N*-back task, calculated as the average of 0- and 2-back task, important for working memory performance; ii) a relational task central for planning and reasoning; iii) a sleep index averaging hours of sleep the month before the fMRI scan. Each subject had a single behavioral score for each of the three behavioral domains. We therefore averaged *PC*_*norm*_ across all nodes in the network to yield a single *PC*_*norm*_ value, for each subject. We show that, using the same bootstrapping procedure outlined previously, mean *PC*_*norm*_ associated with functional brain networks was correlated more strongly with the *N*-back task compared to *PC*, but not the sleep index and relational task (Fig. 7). This suggests that working memory performance relates to the extent of inter-modular connectivity, rather than the size, of modules.

**Figure 7:**
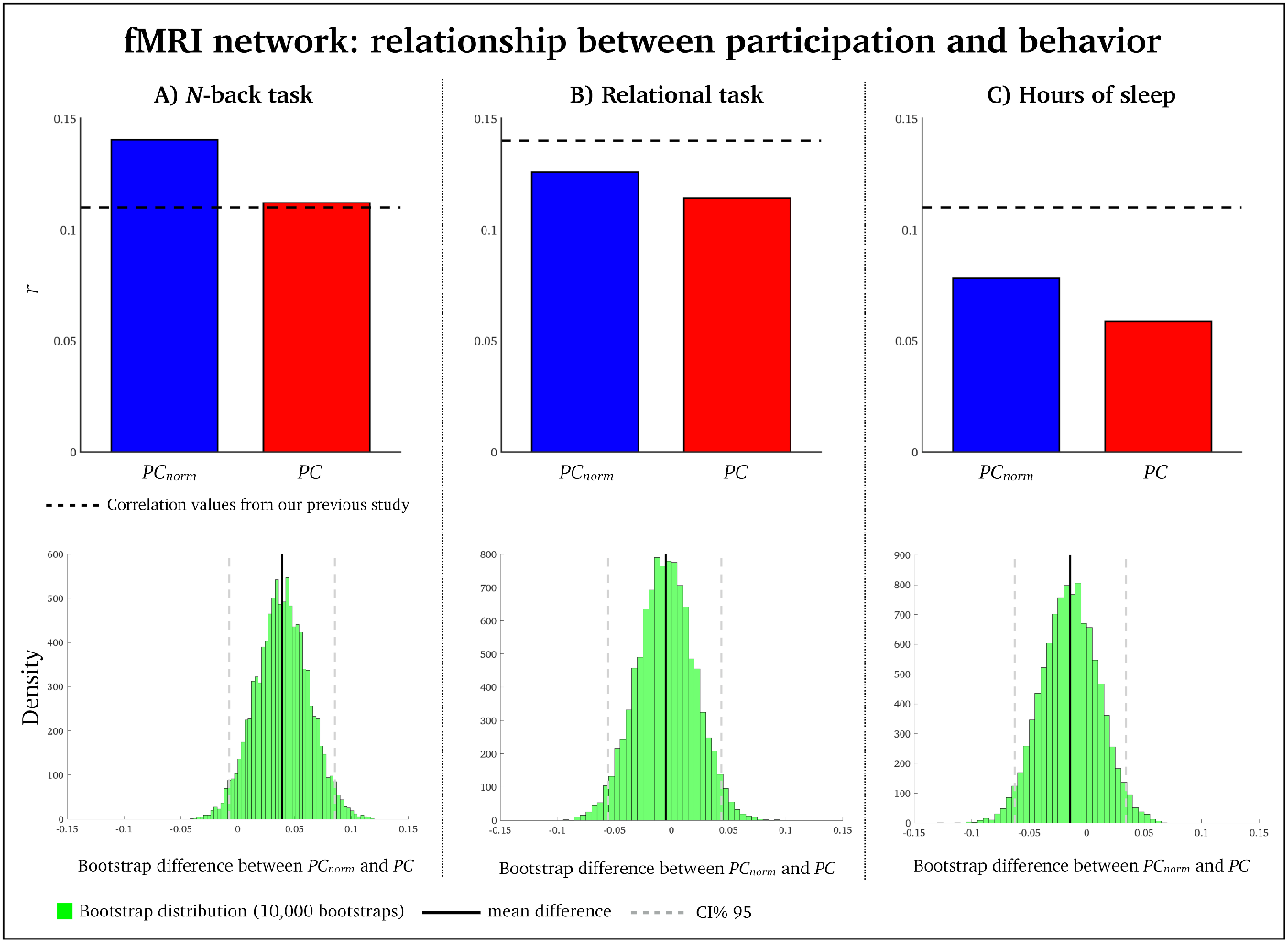
Correlation between *PC*_*norm*_, *PC* and behavior: Inter-individual variation in *PC*_*norm*_ (blue) and *PC* (red) was associated with three behavioral measures (A) *N*-back task, (B) sleep index and (C) relational task. In each panel, the dashed line represents the correlation between modular network switching and behavior obtained in our previous study [8]. Pearson’s *r* between *N*-back task and *PCnorm* was 0.14 (df = 1001, *p* < 0.00001); *r* between *N*-back task and *PC* was 0.11 (df = 1001, *p* < 0.001). Bootstrap difference between *PC*_*norm*_ and *PC* for *N*-back task was 0.04 [95% CI = −0.01 - 0.08]. *r* between sleep and *PC*_*norm*_ was 0.12 (df = 1001, *p* < 0.0001); *r* between sleep and *PC* was 0.13 (df = 1001, *p* < 0.0001). Bootstrap difference between *PC*_*norm*_ and *PC* for sleep was 0.00 [95% CI = −0.05 - 0.05]. *r* between relational task and *PC*_*norm*_ was 0.06 (df = 1001, *p* = 0.07); *r* between relational task and *PC* was 0.07 (df = 1001, *p* = 0.04). Bootstrap difference between *PC*_*norm*_ and *PC* for relational task was 0.01 [95% CI = −0.04 - 0.05].

### *PC*_*norm*_, *PC* and within-module degree *z*-score in real-world networks

*PC* is often interpreted alongside within-module degree *z*-score, a metric that quantifies the normalized degree of intra-modular nodes. First of all, we observed almost no correlation between within-module degree *z*-score and module size (Pearson’s *r*^2^ = 0.01, *γ* = 1). We next projected corresponding values between *PC* and within-module degree *z*-score in a joint histogram which categorizes the intra- and inter-modular statuses of nodes (see Fig 8). Similarly to previous work, most network nodes are peripheral nodes classified as nodes that mainly connect to a single module [6]. However, our joint histogram analysis showed that *PC*_*norm*_ detected more non-hub connector nodes than *PC* in fMRI networks (Fig. 8A). Non-hub connector nodes are classified as nodes that connect to a variety of modules. Airport networks also showed increased *PC*_*norm*_, compared to *PC*, but remained as peripheral nodes (Fig. 8B). On the other hand, *PC* and *PC*_*norm*_ yielded comparable inference about the *C.elegans* network, which primarily comprised peripheral nodes (see Fig. 8C). This suggests that *PC*_*norm*_ may alter the topological structure, and interpretation, of networks where the module size bias is prominent.

**Figure 8:**
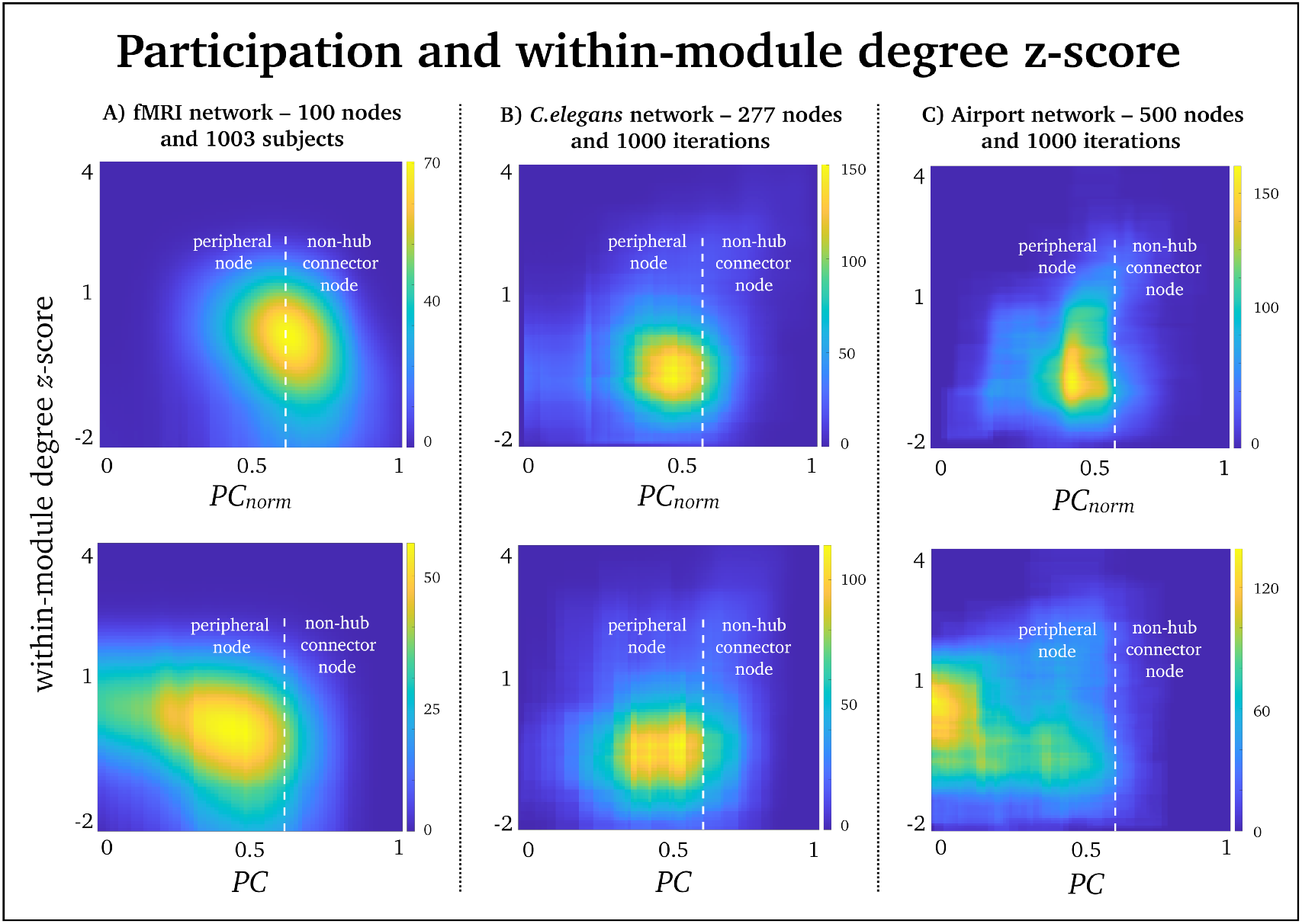
Participation and within-module degree *z*-score: Joint histograms containing the number of simultaneous occurrences between *PCnorm* (top row), *PC* (bottom row) and within-module degree *z*-score in (A) fMRI networks, (B) *C.elegans* networks and (C) airport networks. Within-module degree *z*-score is shown on the vertical axis, while *PC* and *PC*_*norm*_ is shown on the horizontal axis. In accordance with [6], *PC* of 0.62 was used to distinguish between peripheral nodes (*PC* < 0.62) and non-hub connector nodes (*PC* > 0.62).

## Discussion

We proposed a normalized participation coeffcient (*PC*_*norm*_) to alleviate the module size bias inherent to the established formulation of *PC* developed by Guimerà and Amaral [4]. Using brain, *C.elegans*, airport and simulated networks, we demonstrated that our proposed measure of participation: i) alleviates module size bias to a significant extent; ii) preserves the conceptual and mathematical properties of the classic formulation of *PC*; and, iii) yields new insights into the role and function of nodes in the context of their participation across modules, thereby enabling clearer and more intuitive conclusions to be drawn about the role of nodes. Our work should be conceptualized as an extension to Guimerà and Amaral’s classic formulation of *PC* [4], rather than a new measure to include in the network scientist’s toolbox. We advocate use of *PC*_*norm*_ in favour of *PC*, particularly for networks with significant variation in module size. These are the networks for which the module size bias is most detrimental to inference performed based on the classic participation coefficient.

It is important to clarify and highlight the crux of *PC*’s module size bias, namely network modularity. Modularity decomposition methods are designed to search for a collection of nodes that have stronger and more widespread inter-connectedness than we expect by chance [21]. Then, a sizeable collection of inter-connected nodes forming a large module will, more often than not, have low *PC* values without necessarily reflecting the node’s inter-connectedness with other modules. An example of this is the case of airport networks, which showed the strongest module size bias. We observed maximal *PC* in a subset of airports located in the smallest module comprising Central- and South America, that consequently led to high *PC* values (Fig. 6). After correcting this module size bias, *PC*_*norm*_ suggested that the airports with strongest inter-modular connectivity were located in metropolitan cities such as New York, London and Paris where over one third of flights were cross-continental. It is also important to note that *PC*_*norm*_, like *PC*, appears independent from node degree. This is evident as the two airports with highest *PC*_*norm*_ had highly divergent node degree: i) New York had a total of 218 connections (127 intra-modular connections); and ii) Punta Cana in the Dominican Republic had only 52 connections (34 intra-modular connections). Although Punta Cana is not a densely connected airport, it belongs to the large module encompassing North America. This airport has a substantial number of cross-continental flights, mainly to South America, again demonstrating how *PC* may underestimates the integrative nature of nodes within large modules. A specific example where *PC* was less affected by module size was the RMGR neuron in the *C.elegans* nematode. The RMGR neuron of the *C.elegans* nematode, which showed similar values between *PC* and *PC*_*norm*_ (Supplementary Table 2), despite belonging to a spatially large module. This finding may be due to the high fidelity of the *C.elgegans* connectome, which was mapped using electron microscopy, and also the inherent function of this neuron. RMGR is an interneuron located at the head of the *C.elegans* nematode, and it makes gap junctions to multiple other neuron classes [29]. Its main function is to integrate information between extrinsic and intrinsic brain activity that leads to behavioral responses [30]. The function of RGMR therefore provides a plausible explanation why it connects to several modules.

An example of the utility of *PC*_*norm*_ in real-world networks was seen in fMRI brain networks, where we observed increased inter-subject correlation with the *N*-back task, compared to *PC* (Fig. 7A). This task is important for working memory performance [31], a higher-order cognitive process that engages a distributed brain network comprising symmetric and bilateral frontal and parietal cortices [32–34]. We found that *PC*_*norm*_ was relatively high in these regions compared to other areas of the cortex, but this was not as prominent as *PC*. Several reports also suggest that *PC* is correlated with working memory performance [5, 11, 35, 36]. For example, Shine et al. [11] reported increased *PC* in fronto-parietal networks during the *N*-back task, compared to the resting state. This suggests that the human brain may allocate more inter-modular connectivity to meet demands of this cognitively strenuous task. Other brain regions that expressed relatively high values of *PC*_*norm*_ were located in the cerebellum, mainly sub-region Crus I and Lobule VI (see Fig. 4A). These cerebellar sub-regions are also involved in working memory [37–39]. They display connectivity with frontal and parietal cortices [40, 41], and have preferentially expanded over recent evolutionary time [42].

Unlike *PC*, nodes that only have intra-modular connections will not have a *PC*_*norm*_ value of zero. This is because our randomization approach always returns a ‘random’ number to subtract from the original *PC* algorithm. Nevertheless, our simulations show that *PC*_*norm*_ return values close to zero for nodes that only have intra-modular connections, unless these are high-degree nodes (see upper left corner of Fig. 2C). We also found that, in real-world networks, *PC*_*norm*_ remains low when *PC* is zero (Supplementary Fig. 6). An alternative is to conduct a post-hoc correction setting *PC*_*norm*_ nodes to zero if corresponding *PC* values are zero.

We have released a user-friendly MATLAB code for calculation of *PC*_*norm*_, available at GitHub. The current version of *PC*_*norm*_ works for undirected/directed and weighted/binary networks. A limitation of *PC*_*norm*_ is the computational time needed for network randomizations, which can be particularly time consuming for large networks (i.e. > 1000 nodes). To circumvent this issue, we added a parallel computing option in our code that allows for faster computation time, ensuring this process is tractable for all but the largest networks. Nevertheless, as network randomization procedures are becoming increasingly popular, it is imperative to continue optimizing computational capabilities when randomizing edges in large and complex networks.

## Methods

### Modular decomposition of networks

Before estimating *PC* and *PC*_*norm*_ (defined in the Results section - Eqs. 1 and 2), we parsed fMRI, *C.elegans* and airport networks into modules using a Louvain modularity optimization procedure [21] written as:

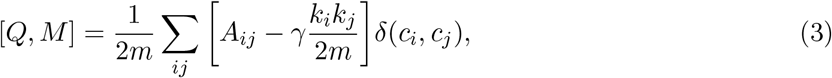

where *A*_*ij*_ represents the binary connectivity edge between nodes *i* and *j*. Louvain modularity optimisation procedure finds nodes with greater intra-modular connectivity that is expected by chance using a null model where *k_i_k_j_* (the sum of the weights of the edges attached to nodes *i* and *j*) is divided by 2*m* (the sum of all of the edge weights in the graph). *c_i_* and *c_j_* are the modules associated with node *i* and *j*, and the module vector output is *M*.

*γ* controls the spatial resolution of modules. A low *γ* (*γ* < 1.5) results in a few large modules, whereas a high *γ* (*γ* > 1.5) returns numerous small modules [20]. We validated our findings across a variety of *γ* parameters between 1 and 2 in 0.1 increments, [43]. The Louvain clustering method also contain heuristics that may cause run-to-run variability due to the degeneracy issue in complex network [44]. This provided us with a good opportunity to assess *PC* across a range of modular structures in the *C.elegans* and airport network where we only had a single network available for analysis. In these two networks, we obtained 1000 different modular outputs. In fMRI networks we computed a single modular output of each of the 1003 subjects using a consensus clustering approach with 1000 iterations. For each modularity run in *C.elegans* and airport networks, and fMRI network subject, we obtained a 1 × *N* vector where *N* is total number of nodes (i.e., 100 nodes for fMRI networks, 277 nodes for the C.elegans neuronal system and 500 nodes for the airports network) comprising *m*-numbers of modules (*M* = all modules).

### Within-module degree *z*-score

To compare *PC* and *PC*_*norm*_ with intra-modular connectivity we computed *within-module degree z-score* [4], i.e., the normalized connectivity of a node within its own module [4]. It is given by:

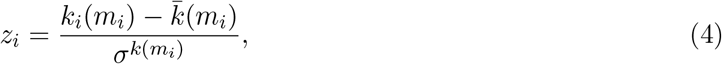

where *m*_*i*_ denotes the module containing *i*; *k*_*i*_(*m*) is the overall degree of node *i* in module *m* (where node *i* is located). 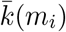 is the mean degree of all nodes in module *m* and 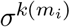 is the standard deviation of all nodes in module *m*.

### Dataset 1: fMRI network

We used 57.6 min (4800 time-points × 0.72s repetition time) of resting state fMRI data from 1003 healthy adults between ages of 22 and 35 years, from the Human Connectome Project [16], that we filtered between frequencies of 0.01 and 0.1 Hz. The fMRI data was distortion, motion, and field bias corrected and normalized into a common Montreal Neurological Institute space. An independent components analysis was conducted to parcellate the brain into 100 regions of interests (55 sub-cortical and 45 cortical brain regions). At the individual level, we generated an undirected binary brain network with 100 nodes using pair-wise Pearson correlation analysis of subject-specific fMRI time series over 100 brain regions. For our main analysis we thresholded networks at a sparsity level 20% (990 edges). We also derived networks at additional sparsity levels of 10% (495 edges), 30% (1485 edges) and 40% (1980 edges – Supplementary Fig. 4)

### Dataset 2: *C.elegans* network

We analyzed two-dimensional spatial representations of the global neuronal network (277 neurons, or nodes, and 2105 connections, or edges) of the nematode *C.elegans*, previously identified by electron microscope reconstructions [18]. This network is directed and binary – i.e., it contains both incoming and outgoing neural connections between brain areas of the *C.elegans*. In this study we only used outgoing connections.

### Dataset 3: Airport network

Airport network was constructed based on flights between the 500 busiest airports (nodes) in the world. Edges were based on the total passenger volume between airports. In this network there were a total of 24009 flights (edges). The existence of flight connections between airports is based on flights within one year from 1 July 2007 to 30 June 2008 [19]. An edge means that there is at least one flight between two airports. This network is directed and binary, meaning it contains incoming and outgoing flights connections between airports. In this study we only used outgoing connections.

## Acknowledgements

This study was supported by the National Health and Medical Research Council (NHMRC) of Australia (no 628952). The Florey Institute of Neuroscience and Mental Health acknowledges the strong support from the Victorian Government and in particular the funding from the Operational Infrastructure Support Grant. We also acknowledge the facilities, and the scientific and technical assistance of the National Imaging Facility (NIF) at the Florey node and The Victorian Biomedical Imaging Capability (VBIC). GJ is supported by an NHMRC practitioner’s fellowship (no 1060312).

The primary fMRI data in this study was provided by the Human Connectome Project, WUMinn Consortium (1U54MH091657; Principal Investigators: David Van Essen and Kamil Ugurbil) funded by the 16 National Institutes of Health (NIH) institutes and centers that support the NIH Blueprint for Neuroscience Research; and by the McDonnell Center for Systems Neuroscience at Washington University. We also thank Marcus Kaiser for releasing network datasets including *C.elegans* and airport networks.

## Author contributions

M.P., A.O., J.M.S., G.D.J., and A.Z. designed research; M.P., A.O., J.M.S., A.Z. performed research; M.P., A.O., analyzed data; and M.P., A.O., J.M.S., G.D.J., and A.Z. wrote the paper.

## Data availability

All fMRI network data can be downloaded from: https://www.humanconnectome.org.

*C.elegans* and airport network data can be downloaded from https://www.dynamic-connectome.org.

*PC*_*norm*_ can be downloaded from https://github.com/omidvarnia/Dynamic_brain_connectivity_analysis.

## Footnotes

1 Q-scores range between 0 and 1 where values proximate to 1 indicate that the network is highly modular, with a predominance of intra-modular connections.

