## Supplementary Figure for "Reducing module size bias of participation coefficient"

### Supplementary materials

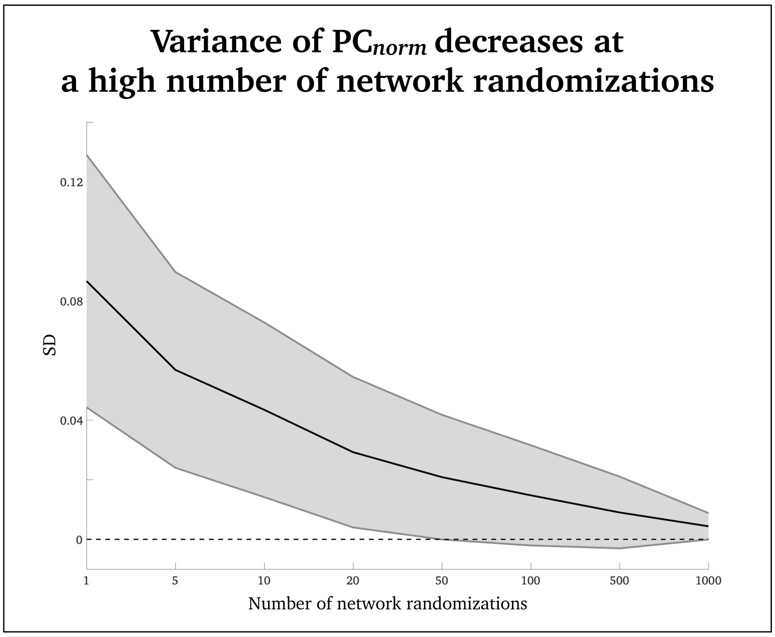


**Figure S1:** We wanted to test how many network randomizations are needed to return a stable estimate of $PC_{norm}$. For this purpose, we used a single randomly selected fMRI networks and derived $PC_{norm}$ from 8 different network randomization parameters (1, 5, 10, 20, 50, 100, 500 and 1000 randomizations). For each of these 8 randomization parameters, we computed 1000 networks. Then the SD across the 1000 networks was computed, for each randomization parameter. We observed less variance across 1000 network iterations at a high number of network randomizations (up to 1000 network randomizations). Nevertheless, variance was also relatively low when computing 100 network randomizations, which may be more feasible in large networks due to the computational burden of network randomizations.

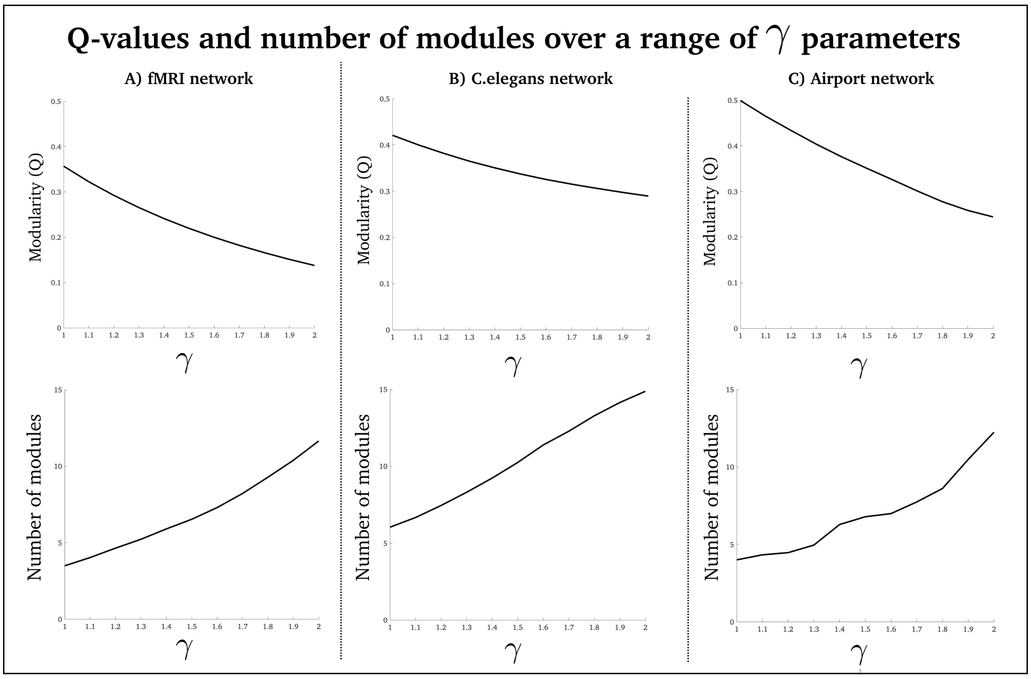


**Figure S2:** We computed modularity (Q-value) the average number of modules across a range of $\gamma$ parameters (1 to 2, in 0.1 increments). In this figure we present the median value across subjects (fMRI – A) and modularity runs (*C.elegans* – B; airport – C). We observed that modularity decreased, and number of modules increased, as $\gamma$ increased, across all three network types. This suggest that the networks may lose their inherent modular structure at high $\gamma$ values.

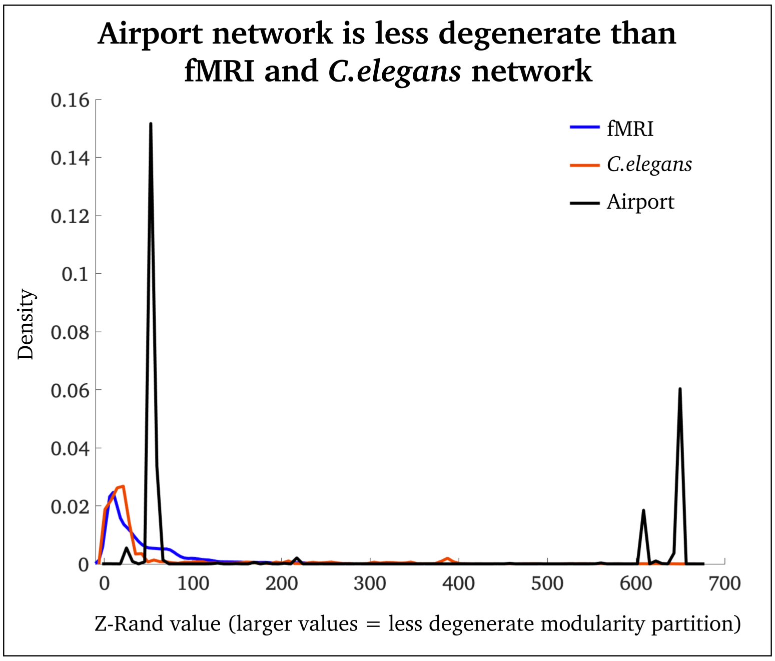


**Figure S3:** In this figure we show that the airport network (black distribution) is less degenerate than fMRI (blue distribution) and *C.elegans* (red distribution) networks, where larger Z-rand values means that the network is less degenerate. Degeneracy refers to the stability of network outputs modularity iterations (less stable outputs = more degeneracy). For a more thorough introduction to the Z-rand metric, refer to Traud et al, SIAM Review 53, 526 (2011).

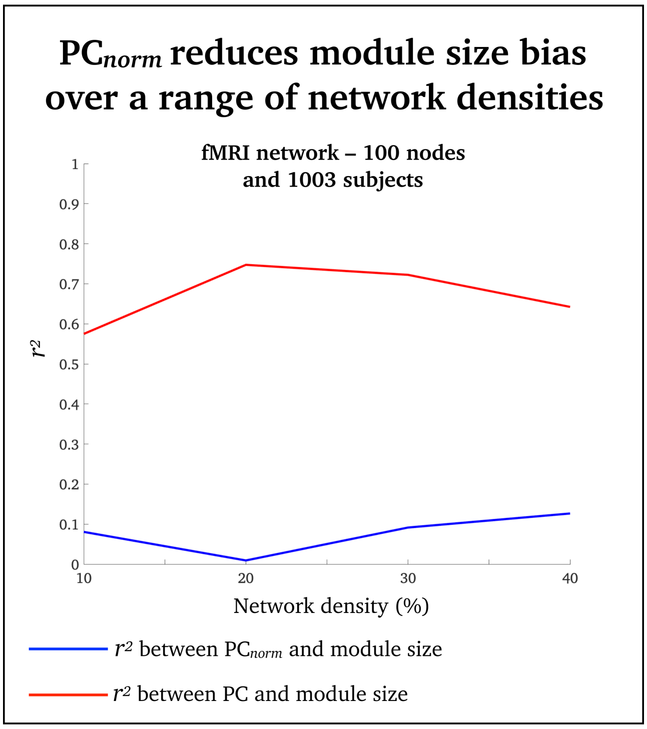


**Figure S4:** $PC_{norm}$ reduces $PC$'s module size bias in fMRI networks, across multiple connection densities Connection density can markedly impact a network's modular architecture as well as other topological properties [23]. Hence, we aimed to test whether variance explained between $PC$, $PC_{norm}$ and module size differed across network densities. To this end, we thresholded each of the 1,003 brain networks to achieve connection densities of 10%, 20%, 30% and 40%. We were not able to use *C.elegans* and airport networks for this analysis as these network datasets were pre-thresholded and binarized before public release. We observed lower correlations between $PC_{norm}$ and module size (Pearson's $r^2$ ranged between 0.01 and 0.13) compared to $PC$ and module size (Pearson's $r^2$ ranged between 0.56 and 0.74), across all connection density thresholds. The bootstrapped $r^2$ difference between $PC_{norm}$ and $PC$ ranged between 0.47 [95% CI = 0.38 – 0.54] and 0.73 [95% CI = 0.64 – 0.76], across all connection density thresholds (Supplementary figure 3).

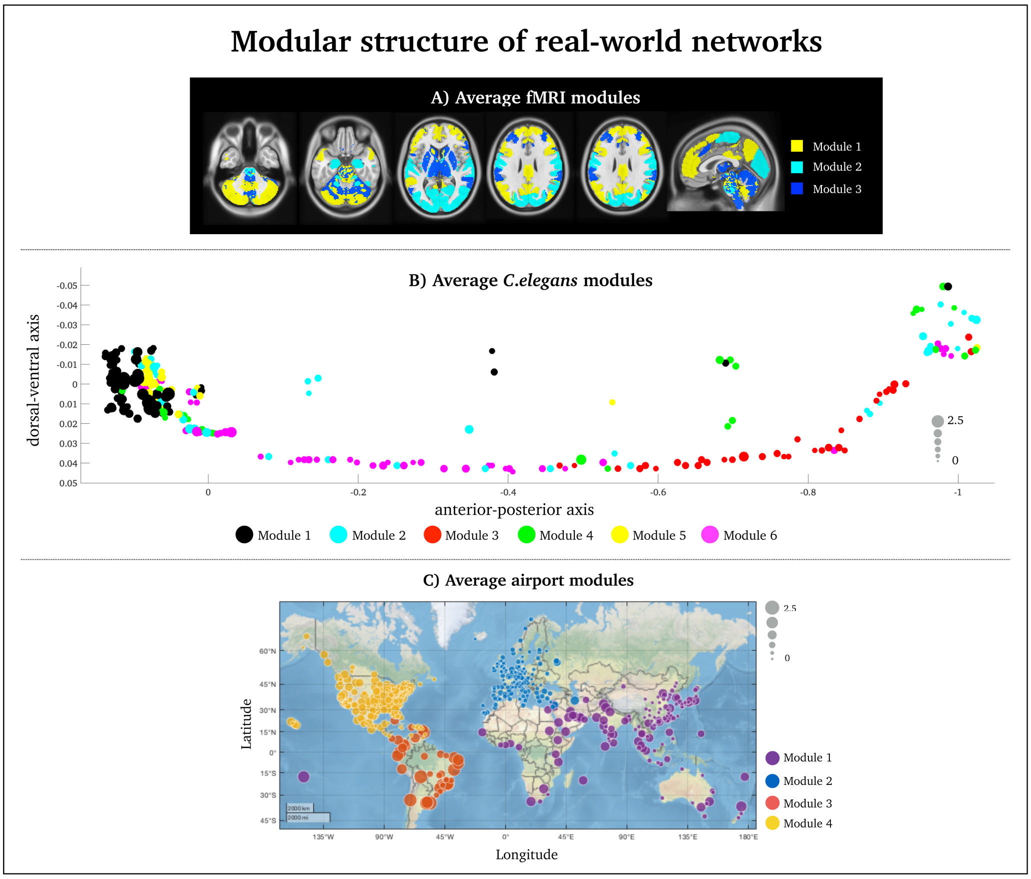


**Figure S5:** Modularity structure in real-world networks. (A) Group-level fMRI modularity map derived from a consensus clustering algorithm. Consensus clustering algorithms derive a single fMRI module ($1 \times 100$ node vector) that best describes the modular information from all subjects and nodes ($1003$ subjects $\times$ $100$ node array) [43]. (B) The colors denote distinct modules derived from a consensus clustering algorithm across 1000 modularity iterations. The black module displayed on the left of the figure comprises inhibitory GABAergic neurons that belong to the largest module of the nematode encompassing RMDDL, RICR, RMDVL, RMDDR and RMDVR neurons. (C) Colors denote modules derived from a consensus clustering algorithm across 1000 Louvain modularity iterations.

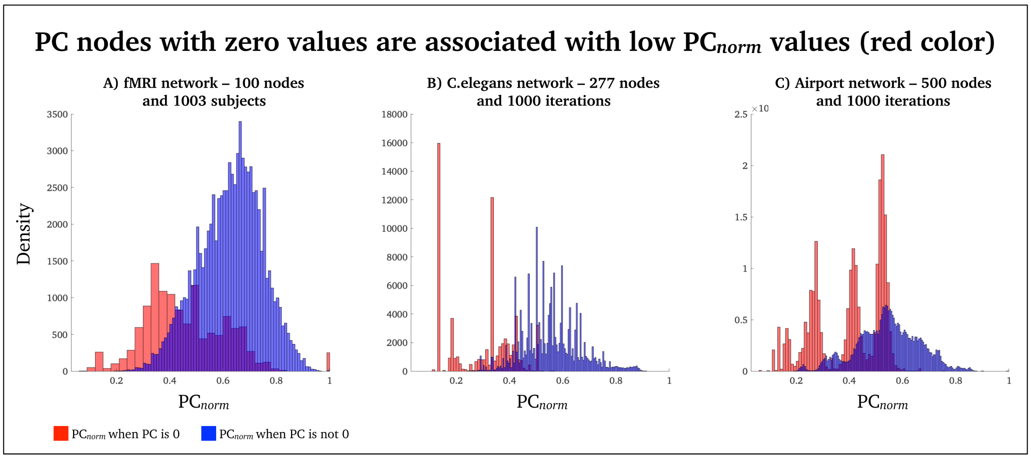


**Figure S6:** As we discussed in the main manuscript, nodes that only have intra-modular connections will not have a $PC_{norm}$ value of zero (unlike $PC$). This is because our randomization approach always generates a 'random' number to subtract from the original $PC$ algorithm. The red distributions in this figure reflects all $PC_{norm}$ nodes that corresponds to $PC$ nodes with values of zero. It is evident that $PC_{norm}$ remains low when $PC$ is zero (red distribution) compared to when $PC$ is not zeros (blue distribution), across A) fMRI, B) *C.elegans* and C) airport networks.

***Supplementary table 1: fMRI* network:** Full set of nodal results for $PC_{norm}$ and PC, and z-score differences between $PC_{norm}$ and PC. Note we used a z-score since $PC_{norm}$ tends to yield higher values than PC -- see Fig. 2 and 3). The last column displays the module community affiliation for each node.

| | fMRI node (AAL definition) | $PC_{norm}$ | PC | Z-score difference | Module |
|---|---|---|---|---|---|
| 1 | Cerebelum_6_L | 0.725951212 | 0.489386403 | 0.567616364 | 1 |
| 2 | Cerebelum_Crus1_BIL | 0.719282621 | 0.456551875 | 1.093689928 | 2 |
| 3 | Parietal_Inf_R | 0.708799866 | 0.499882327 | 0.011760156 | 1 |
| 4 | Middle Frontal Gyrus_BIL | 0.707511318 | 0.521348924 | -0.445738491 | 1 |
| 5 | Precuneus_BIL | 0.701201163 | 0.53041552 | -0.75489244 | 3 |
| 6 | Cerebelum_Crus1_R | 0.699871394 | 0.428270238 | 1.272032053 | 2 |
| 7 | Insula_R | 0.696412289 | 0.520064242 | -0.643058737 | 2 |
| 8 | Parietal_Inf_R | 0.690456671 | 0.482024811 | 0.001995438 | 1 |
| 9 | Cerebelum_Crus1_R | 0.689182966 | 0.418089464 | 1.261825526 | 2 |
| 10 | Cerebelum_Crus2_BIL | 0.687837526 | 0.402389567 | 1.550425938 | 2 |
| 11 | Cerebelum_Crus2_L | 0.685835304 | 0.407177106 | 1.413915897 | 1 |
| 12 | Cerebelum_Crus1_L | 0.683617476 | 0.445404201 | 0.600759243 | 2 |
| 13 | Caudate_dorsal_BIL | 0.683271034 | 0.460794317 | 0.284371278 | 1 |
| 14 | Cerebelum_6_BIL | 0.68268362 | 0.437801579 | 0.734836656 | 2 |
| 15 | Cerebelum_8_R | 0.680244395 | 0.41485296 | 1.147183857 | 2 |
| 16 | Cerebelum_7b_R | 0.679796826 | 0.401726195 | 1.402102678 | 1 |
| 17 | Precuneus_BIL | 0.679376428 | 0.483770649 | -0.255876548 | 1 |
| 18 | Parietal_Sup_L | 0.67933796 | 0.514261427 | -0.869675716 | 1 |
| 19 | Thalamus_L | 0.678833896 | 0.443459874 | 0.543675239 | 1 |
| 20 | Cerebelum_Crus1_R | 0.672781211 | 0.392688835 | 1.442750444 | 1 |
| 21 | Insula_L | 0.668925572 | 0.509384743 | -0.980972612 | 1 |
| 22 | Cerebelum_8_L | 0.663320703 | 0.412712444 | 0.84996391 | 2 |
| 23 | Cerebelum_Crus2_L | 0.660374916 | 0.379925207 | 1.449934721 | 1 |
| 24 | Cerebelum_6_L | 0.658623439 | 0.379776311 | 1.417714377 | 2 |
| 25 | Cerebelum_7b_L | 0.657134246 | 0.396904637 | 1.043403875 | 1 |
| 26 | Vermis_8 | 0.65628065 | 0.414777894 | 0.666895176 | 3 |
| 27 | Vermis_4 | 0.648899941 | 0.415845938 | 0.497030613 | 1 |
| 28 | Cerebelum_6_L | 0.646939909 | 0.373842709 | 1.302110443 | 1 |
| 29 | Cerebelum_6_R | 0.643372756 | 0.362136018 | 1.465758171 | 2 |
| 30 | Cerebelum_6_R | 0.643307166 | 0.380369987 | 1.097840336 | 1 |
| 31 | Inferior Parietal Lobule_R | 0.642520319 | 0.434530476 | -0.00689144 | 3 |
| 32 | Cerebelum_Crus2_R | 0.642511289 | 0.356204154 | 1.567699938 | 1 |
| 33 | Middle Frontal Gyrus_R | 0.642210271 | 0.472533118 | -0.777178945 | 2 |
| 34 | Cerebelum_8_R | 0.641895261 | 0.379818496 | 1.080541471 | 1 |
| 35 | Occipital_Mid_BIL | 0.640190742 | 0.464514998 | -0.656575566 | 1 |
| 36 | Cerebelum_Crus1_L | 0.638851414 | 0.349244781 | 1.634037279 | 2 |
| 37 | Cerebelum_8_L | 0.637358197 | 0.433325552 | -0.086452018 | 2 |
| 38 | Cerebelum_6_L | 0.635117564 | 0.381186409 | 0.91677167 | 1 |
| 39 | Cerebelum_Crus1_R | 0.633504629 | 0.352896195 | 1.453125934 | 1 |
| 40 | Cerebelum_7b_R | 0.630269707 | 0.351818813 | 1.409747967 | 2 |
| 41 | Caudate_ventral_BIL | 0.629419965 | 0.39964153 | 0.431174388 | 1 |
| 42 | Superior Frontal Gyrus_R | 0.626212259 | 0.361951022 | 1.124460893 | 2 |
| 43 | Angular_L | 0.624061779 | 0.393933949 | 0.438199068 | 1 |
| 44 | Occipital_Mid_R | 0.623814323 | 0.470296181 | -1.10206044 | 2 |
| 45 | Occipital_Sup_R | 0.623754267 | 0.458167468 | -0.859416657 | 1 |
| 46 | Middle Temporal Gyrus_BIL | 0.619242895 | 0.442314584 | -0.631392361 | 2 |
| 47 | Frontal_Mid_Orb_L | 0.617835871 | 0.398686201 | 0.217480052 | 1 |
| 48 | Cerebelum_Crus1_L | 0.617033927 | 0.313827139 | 1.907472245 | 2 |
| 49 | Cerebelum_6_R | 0.617027459 | 0.391339691 | 0.348930366 | 2 |
| 50 | Cerebelum_8_L | 0.616260913 | 0.404553054 | 0.067860315 | 2 |
| 51 | Putamen_ant_BIL | 0.615960006 | 0.397418914 | 0.205244413 | 2 |
| 52 | Putamen_post_BIL | 0.615396757 | 0.384433481 | 0.454995947 | 2 |
| 53 | Cerebelum_6_L | 0.612024349 | 0.372582107 | 0.62546797 | 2 |
| 54 | Putamen_mid_BIL | 0.60662399 | 0.404821032 | -0.131280519 | 2 |
| 55 | Cerebelum_8_R | 0.606290296 | 0.402041408 | -0.082104406 | 2 |
| 56 | Frontal_Sup_Medial_R | 0.606018819 | 0.360412077 | 0.749406974 | 2 |
| 57 | Angular_L | 0.601316248 | 0.342745353 | 1.010054962 | 2 |
| 58 | Thalamus_BIL | 0.600835109 | 0.41521247 | -0.456590426 | 3 |
| 59 | Frontal_Inf_Orb_L | 0.598774969 | 0.392050394 | -0.032330017 | 2 |
| 60 | Cerebelum_Crus1_R | 0.596773279 | 0.300963207 | 1.758759188 | 2 |
| 61 | Parietal_Inf_R | 0.596010994 | 0.337935742 | 1.000089929 | 2 |
| 62 | Parietal_Inf_L | 0.593605293 | 0.450380407 | -1.309009282 | 1 |
| 63 | Medulla | 0.589977118 | 0.383980779 | -0.046971401 | 3 |
| 64 | Brainstem_L | 0.586985607 | 0.36197052 | 0.33540592 | 2 |
| 65 | Calcarine_L | 0.581827878 | 0.435031595 | -1.237205304 | 3 |
| 66 | Temporal_Mid_L | 0.581701092 | 0.40689825 | -0.6741255 | 2 |
| 67 | Occipital_Mid_L | 0.578437473 | 0.415595828 | -0.914608752 | 3 |
| 68 | Parietal_Sup_L | 0.570980573 | 0.389978683 | -0.549491883 | 2 |
| 69 | Putamen_L | 0.56969866 | 0.381170914 | -0.398182425 | 3 |

| | | | | | |
|---|---|---|---|---|---|
| 70 | Substania Nigra_L | 0.566440702 | 0.391396332 | -0.669269517 | 1 |
| 71 | Parahippocampa Gyrus_BIL | 0.565564426 | 0.419427381 | -1.250459483 | 2 |
| 72 | Hippocampus_R | 0.563331005 | 0.373402657 | -0.370022936 | 3 |
| 73 | Substania Nigra_R | 0.56141693 | 0.367824025 | -0.29634596 | 2 |
| 74 | Pons | 0.560676466 | 0.411730107 | -1.193977432 | 2 |
| 75 | Brainstem | 0.560323184 | 0.398835372 | -0.941827964 | 1 |
| 76 | Precuneus_L | 0.556986968 | 0.369647092 | -0.42206488 | 3 |
| 77 | Right Brainstem | 0.555267266 | 0.412840965 | -1.325065044 | 1 |
| 78 | Occipital_Mid_L | 0.555239553 | 0.392892792 | -0.924558558 | 2 |
| 79 | Occipital_Inf_L | 0.545434379 | 0.381949455 | -0.90167545 | 3 |
| 80 | Midbrain | 0.53727558 | 0.399713988 | -1.4228714 | 2 |
| 81 | Postcentral_L | 0.52553897 | 0.332992653 | -0.317387908 | 1 |
| 82 | Occipital_Sup_R | 0.522501286 | 0.375968499 | -1.242502982 | 1 |
| 83 | Precentral_R | 0.521512829 | 0.337349962 | -0.485939556 | 2 |
| 84 | Midbrain | 0.51916125 | 0.454768214 | -1.393945971 | 1 |
| 85 | SupraMarginal_R | 0.518996924 | 0.319905905 | -0.185804828 | 1 |
| 86 | Parietal Inf_R | 0.513079098 | 0.33479285 | -0.604090662 | 2 |
| 87 | Thalamus_R | 0.509204671 | 0.376659107 | -1.523720063 | 3 |
| 88 | Lingual_BIL | 0.505338366 | 0.319742756 | -0.457133845 | 1 |
| 89 | Paracentral_Lobule_BIL | 0.503114286 | 0.350230544 | -1.114815231 | 1 |
| 90 | Postcentral_BIL | 0.503097943 | 0.327131223 | -0.650725402 | 2 |
| 91 | Temporal_Mid_BIL | 0.501047122 | 0.337169126 | -0.893772636 | 2 |
| 92 | Pons | 0.500379445 | 0.403477024 | -1.240335576 | 1 |
| 93 | Occipital_Sup_R | 0.499012248 | 0.312912064 | -0.446989249 | 2 |
| 94 | Postcentral_R | 0.496588428 | 0.316175691 | -0.561336984 | 2 |
| 95 | Occipital_Mid_R | 0.496150775 | 0.311384889 | -0.473815674 | 2 |
| 96 | Temporal_Sup_BIL | 0.484510726 | 0.307651196 | -0.632775203 | 1 |
| 97 | Occipital_Sup_BIL | 0.481284986 | 0.306194976 | -0.668351912 | 1 |
| 98 | Occipital_Sup_L | 0.467186012 | 0.29239596 | -0.674382642 | 1 |
| 99 | Fusiform_BIL | 0.457127764 | 0.258010197 | -0.18527107 | 2 |
| 100 | SupraMarginal_L | 0.454337824 | 0.321235356 | -0.342712047 | 1 |

***Supplementary table 2: C.elegans* network:** Full set of nodal results for $PC_{norm}$ and PC, and z-score differences between $PC_{norm}$ and PC. Note we used a z-score since $PC_{norm}$ tends to yield higher values than PC -- see Fig. 2 and 3). The last column displays the module community affiliation for each node. See https://www.wormatlas.org/neurons/Individual%20Neurons/Neuronframeset.html for full naming, definition and function of specific *C.elegans* neurons.

| | Node | $PC_{norm}$ | PC | Z-score difference | Module |
|---|---|---|---|---|---|
| 1 | AVER | 0.87188934 | 0.754524082 | 0.803904556 | 3 |
| 2 | AVAR | 0.850417827 | 0.751160024 | 0.648866855 | 3 |
| 3 | AVAL | 0.841479159 | 0.753069215 | 0.55598649 | 4 |
| 4 | AVBR | 0.832115473 | 0.744365741 | 0.550333698 | 4 |
| 5 | AVBL | 0.800101977 | 0.737799184 | 0.332454664 | 4 |
| 6 | AVEL | 0.799190253 | 0.697926538 | 0.666041661 | 3 |
| 7 | AVDR | 0.745524256 | 0.677048907 | 0.385304649 | 4 |
| 8 | SAADR | 0.726844052 | 0.7234 | -0.171499293 | 5 |
| 9 | AVDL | 0.725415034 | 0.671363905 | 0.261803152 | 4 |
| 10 | RMGR | 0.723789379 | 0.70277551 | -0.021064905 | 5 |
| 11 | RIMR | 0.722040586 | 0.685944637 | 0.108069255 | 5 |
| 12 | PVCL | 0.717077872 | 0.6878912 | 0.048911393 | 4 |
| 13 | PVCR | 0.711504866 | 0.665739645 | 0.190858447 | 4 |
| 14 | RIML | 0.709353064 | 0.688648889 | -0.023716529 | 5 |
| 15 | SMBDR | 0.709172506 | 0.67614876 | 0.081764764 | 5 |
| 16 | RIGL | 0.698700791 | 0.68998 | -0.126319365 | 2 |
| 17 | SMBDL | 0.691542967 | 0.6649375 | 0.026810875 | 5 |
| 18 | RIGR | 0.682752057 | 0.69403125 | -0.297561068 | 2 |
| 19 | RIAR | 0.68186209 | 0.578097633 | 0.687453241 | 5 |
| 20 | RIAL | 0.678463093 | 0.587951389 | 0.573981956 | 5 |
| 21 | AVJL | 0.673746679 | 0.59897 | 0.439257217 | 4 |
| 22 | AIBR | 0.672858478 | 0.6596512 | -0.087905654 | 1 |
| 23 | AVJR | 0.667032021 | 0.605767313 | 0.323566479 | 4 |
| 24 | SAADL | 0.653323653 | 0.706081633 | -0.652706254 | 5 |
| 25 | RIS | 0.647143517 | 0.571755102 | 0.444494964 | 5 |
| 26 | ADER | 0.646803824 | 0.616611111 | 0.057525203 | 4 |
| 27 | RIBL | 0.643620868 | 0.55365625 | 0.569297749 | 2 |
| 28 | RMFL | 0.641424566 | 0.637591837 | -0.168171398 | 5 |
| 29 | DD2 | 0.641409023 | 0.603093333 | 0.127074877 | 3 |
| 30 | DVA | 0.64124183 | 0.653731405 | -0.307924469 | 4 |
| 31 | VB1 | 0.640809319 | 0.72224 | -0.898204556 | 3 |
| 32 | ADEL | 0.631312445 | 0.60216 | 0.048618337 | 5 |
| 33 | AVL | 0.627692227 | 0.54575 | 0.500609304 | 4 |
| 34 | AVKR | 0.62530564 | 0.619263218 | -0.149251809 | 5 |
| 35 | ADAR | 0.623203242 | 0.62355102 | -0.203965293 | 5 |
| 36 | SABD | 0.621880552 | 0.570958678 | 0.235010189 | 3 |
| 37 | SMBVL | 0.617380281 | 0.65644898 | -0.53549737 | 1 |
| 38 | ADAL | 0.612775816 | 0.621888889 | -0.279014544 | 1 |
| 39 | URXL | 0.612358544 | 0.553033033 | 0.306962901 | 2 |
| 40 | RIBR | 0.612235515 | 0.499446192 | 0.764724973 | 5 |
| 41 | AVKL | 0.611602845 | 0.576077775 | 0.10318133 | 5 |
| 42 | BDUR | 0.610350967 | 0.542333333 | 0.381385657 | 4 |
| 43 | BDUL | 0.605318518 | 0.583833834 | -0.01703374 | 4 |
| 44 | DVC | 0.603067698 | 0.5429 | 0.314173784 | 2 |
| 45 | RMHL | 0.603062953 | 0.535341365 | 0.378850876 | 5 |
| 46 | RIPL | 0.602471823 | 0.438602041 | 1.202080562 | 5 |
| 47 | DA8 | 0.601391266 | 0.578760331 | -0.007219429 | 6 |
| 48 | AQR | 0.601340044 | 0.607888889 | -0.257059395 | 4 |
| 49 | RIR | 0.599914812 | 0.612975207 | -0.312811881 | 2 |
| 50 | FLPL | 0.598544558 | 0.548136273 | 0.230612799 | 4 |
| 51 | BAGR | 0.594557278 | 0.597375 | -0.225113178 | 2 |
| 52 | SABVL | 0.593533733 | 0.549111111 | 0.179362994 | 3 |
| 53 | AUAR | 0.592592963 | 0.62944 | -0.516475302 | 5 |
| 54 | OLLR | 0.590665538 | 0.46525 | 0.872831786 | 5 |
| 55 | RMGL | 0.587483553 | 0.468 | 0.822041588 | 5 |
| 56 | RMHR | 0.585681016 | 0.45622582 | 0.907419706 | 5 |
| 57 | AUAL | 0.584712572 | 0.585877551 | -0.210962239 | 2 |
| 58 | SIBVL | 0.580950739 | 0.465611111 | 0.786560917 | 5 |
| 59 | RMEV | 0.578849947 | 0.425003516 | 1.116259707 | 5 |
| 60 | AIZL | 0.574583317 | 0.579289256 | -0.241280263 | 1 |
| 61 | SMBVR | 0.574003944 | 0.607020408 | -0.483677579 | 1 |
| 62 | ALML | 0.573906115 | 0.539070352 | 0.097279431 | 5 |
| 63 | ADLL | 0.573456303 | 0.549777778 | 0.001750134 | 4 |
| 64 | PDEL | 0.571604532 | 0.603985749 | -0.478238541 | 4 |
| 65 | OLQVL | 0.571197681 | 0.4575 | 0.772502418 | 5 |
| 66 | DD1 | 0.568701325 | 0.484648711 | 0.51867863 | 3 |

| | | | | | |
|---|---|---|---|---|---|
| 67 | AS5 | 0.568014997 | 0.449975309 | 0.809679088 | 3 |
| 68 | DB5 | 0.56761642 | 0.538055556 | 0.052115255 | 6 |
| 69 | VD1 | 0.56744537 | 0.483398955 | 0.518625556 | 3 |
| 70 | RIH | 0.567310708 | 0.38504142 | 1.359618818 | 5 |
| 71 | DD4 | 0.564536145 | 0.568319444 | -0.233380539 | 6 |
| 72 | URXR | 0.562172023 | 0.457936281 | 0.691488427 | 5 |
| 73 | AS1 | 0.561330095 | 0.461497155 | 0.653791231 | 3 |
| 74 | DA5 | 0.559985812 | 0.43142 | 0.899804722 | 3 |
| 75 | RMDR | 0.559553639 | 0.339638917 | 1.681942488 | 5 |
| 76 | SABVR | 0.558746515 | 0.50474059 | 0.261416113 | 4 |
| 77 | DD3 | 0.558440096 | 0.48624 | 0.417196279 | 3 |
| 78 | VA12 | 0.558421336 | 0.530868059 | 0.034926115 | 6 |
| 79 | HSNR | 0.557181027 | 0.468773333 | 0.555967219 | 3 |
| 80 | SMDDL | 0.556831482 | 0.344183176 | 1.619726764 | 5 |
| 81 | DVB | 0.553558696 | 0.49321875 | 0.315648585 | 2 |
| 82 | RIPR | 0.552366404 | 0.340485207 | 1.613158708 | 5 |
| 83 | VD4 | 0.551797376 | 0.447641563 | 0.690804075 | 3 |
| 84 | SAAVR | 0.551321787 | 0.491310685 | 0.312832994 | 5 |
| 85 | PVQL | 0.547616604 | 0.445027778 | 0.677387385 | 1 |
| 86 | HSNL | 0.546800579 | 0.460066116 | 0.541640865 | 3 |
| 87 | ASGR | 0.544494291 | 0.582125 | -0.523185169 | 1 |
| 88 | ASHR | 0.542021813 | 0.550428217 | -0.272963991 | 1 |
| 89 | PVPR | 0.540580246 | 0.493234806 | 0.204388427 | 4 |
| 90 | VA6 | 0.540023105 | 0.42489403 | 0.784758149 | 3 |
| 91 | DB3 | 0.539552472 | 0.49930203 | 0.143640402 | 3 |
| 92 | PVPL | 0.539434165 | 0.489567839 | 0.225972486 | 4 |
| 93 | DA6 | 0.53613316 | 0.522400535 | -0.083407585 | 6 |
| 94 | ASKL | 0.535892447 | 0.47230263 | 0.343474281 | 1 |
| 95 | RMED | 0.535103318 | 0.42352 | 0.754399052 | 5 |
| 96 | RIFL | 0.531020525 | 0.52492485 | -0.148795844 | 4 |
| 97 | AWBL | 0.530759057 | 0.555173293 | -0.410024511 | 1 |
| 98 | ADFR | 0.530274365 | 0.499625 | 0.061435097 | 1 |
| 99 | DA2 | 0.530214902 | 0.426519916 | 0.686858422 | 3 |
| 100 | PQR | 0.530142425 | 0.479362912 | 0.233791275 | 4 |
| 101 | AIZR | 0.529796473 | 0.500115702 | 0.053141906 | 1 |
| 102 | PVWL | 0.527370939 | 0.488526077 | 0.131605699 | 4 |
| 103 | SIBVR | 0.525853725 | 0.375 | 1.090635879 | 5 |
| 104 | PVR | 0.525498708 | 0.47437037 | 0.236777947 | 4 |
| 105 | AVHR | 0.523937006 | 0.403024793 | 0.834273903 | 4 |
| 106 | VD3 | 0.518441763 | 0.393358903 | 0.869983373 | 3 |
| 107 | PHBL | 0.517129113 | 0.496849943 | -0.027355456 | 4 |
| 108 | VA11 | 0.516995245 | 0.429906932 | 0.544670561 | 6 |
| 109 | DA9 | 0.515226962 | 0.491970339 | -0.001862233 | 6 |
| 110 | URADL | 0.514979632 | 0.445364619 | 0.39506256 | 5 |
| 111 | URADR | 0.514342 | 0.444444444 | 0.397481713 | 5 |
| 112 | VA10 | 0.514313513 | 0.459166667 | 0.271184787 | 6 |
| 113 | PDER | 0.513137744 | 0.499498998 | -0.084211385 | 4 |
| 114 | VD6 | 0.512309061 | 0.446779853 | 0.360079527 | 3 |
| 115 | PVNR | 0.51048801 | 0.39753125 | 0.766158582 | 4 |
| 116 | RMFR | 0.506886964 | 0.427402863 | 0.479562598 | 5 |
| 117 | RIFR | 0.50632863 | 0.403807615 | 0.676806775 | 4 |
| 118 | VD12 | 0.502386789 | 0.480147796 | -0.010575271 | 6 |
| 119 | AS10 | 0.50131257 | 0.38604345 | 0.785957227 | 6 |
| 120 | CEPDR | 0.496916502 | 0.281685776 | 1.641837686 | 5 |
| 121 | SMDDR | 0.49607419 | 0.330631118 | 1.215551215 | 5 |
| 122 | AIAL | 0.493456509 | 0.411572667 | 0.500109404 | 1 |
| 123 | RICL | 0.493161228 | 0.257777778 | 1.814387165 | 5 |
| 124 | SMDVR | 0.491356161 | 0.249104283 | 1.873195269 | 5 |
| 125 | PVQR | 0.491174759 | 0.348625 | 1.019536559 | 4 |
| 126 | LUAL | 0.489301321 | 0.503755669 | -0.324747033 | 4 |
| 127 | VD5 | 0.487420582 | 0.360851927 | 0.882704882 | 3 |
| 128 | IL1DR | 0.483364612 | 0.32024 | 1.19570035 | 5 |
| 129 | IL1VR | 0.48249764 | 0.278 | 1.549940016 | 5 |
| 130 | IL1DL | 0.481883626 | 0.32088 | 1.17754027 | 5 |
| 131 | VA5 | 0.479996679 | 0.33698092 | 1.023526496 | 3 |
| 132 | PVWR | 0.479617162 | 0.446578631 | 0.081891359 | 4 |
| 133 | IL1VL | 0.479509647 | 0.277777778 | 1.526259236 | 5 |
| 134 | AS2 | 0.479434762 | 0.461061224 | -0.043671663 | 3 |
| 135 | RMDL | 0.477765481 | 0.243496311 | 1.804846595 | 5 |
| 136 | VC2 | 0.477676492 | 0.345485436 | 0.930844401 | 3 |
| 137 | DA1 | 0.4772101 | 0.327350622 | 1.082123042 | 3 |
| 138 | RICR | 0.474975069 | 0.177384615 | 2.347009249 | 5 |
| 139 | DB4 | 0.474442485 | 0.429154775 | 0.186769958 | 3 |
| 140 | ALNL | 0.473633357 | 0.5 | -0.426741205 | 5 |
| 141 | VC5 | 0.471821577 | 0.448022312 | 0.002783909 | 2 |

| | | | | | |
|---|---|---|---|---|---|
| 142 | SIAVR | 0.471641456 | 0.341552941 | 0.912842246 | 5 |
| 143 | VA4 | 0.47062233 | 0.494707151 | -0.407204034 | 3 |
| 144 | RID | 0.470080085 | 0.4485 | -0.016216905 | 4 |
| 145 | AVM | 0.463153439 | 0.515655024 | -0.650510981 | 4 |
| 146 | AS3 | 0.463112168 | 0.425167858 | 0.123895093 | 3 |
| 147 | AS6 | 0.461302801 | 0.474675768 | -0.315488151 | 3 |
| 148 | AS4 | 0.460658081 | 0.425164332 | 0.102913157 | 3 |
| 149 | DD6 | 0.456221029 | 0.402163265 | 0.261859961 | 6 |
| 150 | AWBR | 0.453413318 | 0.469518717 | -0.338883486 | 1 |
| 151 | PVNL | 0.451690583 | 0.297323961 | 1.120713633 | 4 |
| 152 | DA3 | 0.45163069 | 0.310834958 | 1.004518428 | 3 |
| 153 | AIAR | 0.450658601 | 0.368107873 | 0.505819343 | 1 |
| 154 | PVT | 0.449925255 | 0.445553603 | -0.16355709 | 4 |
| 155 | ASHL | 0.449880197 | 0.47341954 | -0.402533602 | 4 |
| 156 | LUAR | 0.447620159 | 0.48102981 | -0.487044084 | 4 |
| 157 | ALNR | 0.445121312 | 0.5 | -0.670863954 | 5 |
| 158 | AS11 | 0.445086132 | 0.443406769 | -0.186608722 | 6 |
| 159 | AVG | 0.44275169 | 0.268336155 | 1.292374265 | 4 |
| 160 | SAAVL | 0.44271288 | 0.395946924 | 0.199426839 | 5 |
| 161 | FLPR | 0.439357681 | 0.463477366 | -0.407502541 | 4 |
| 162 | DA7 | 0.438185466 | 0.478541667 | -0.54652108 | 6 |
| 163 | PHBR | 0.436598306 | 0.444444444 | -0.268166936 | 4 |
| 164 | VD7 | 0.434131652 | 0.377681452 | 0.28234423 | 6 |
| 165 | RIVR | 0.430167614 | 0.338916968 | 0.580308845 | 5 |
| 166 | VD13 | 0.428631317 | 0.483376793 | -0.66972338 | 6 |
| 167 | DB7 | 0.427679625 | 0.461374002 | -0.489481939 | 6 |
| 168 | BAGL | 0.424696177 | 0.5 | -0.845745838 | 2 |
| 169 | RMDDL | 0.424310334 | 0.111774281 | 2.474974843 | 5 |
| 170 | RMDVL | 0.422332494 | 0.133219373 | 2.274425545 | 5 |
| 171 | AVFL | 0.422312095 | 0.29202542 | 0.914538904 | 4 |
| 172 | AVFR | 0.42230374 | 0.287269272 | 0.955189951 | 4 |
| 173 | VD11 | 0.420540594 | 0.339258517 | 0.494957038 | 6 |
| 174 | RMDVR | 0.419663322 | 0.135252214 | 2.234166475 | 5 |
| 175 | RMDDR | 0.418713682 | 0.130752688 | 2.264560922 | 5 |
| 176 | VA2 | 0.417048118 | 0.409914894 | -0.139912257 | 3 |
| 177 | OLLL | 0.414696331 | 0.381831984 | 0.080399982 | 5 |
| 178 | VA3 | 0.414514614 | 0.400199468 | -0.078419985 | 3 |
| 179 | ADLR | 0.414458706 | 0.444444444 | -0.457728232 | 1 |
| 180 | CEPDL | 0.414303209 | 0.281959184 | 0.93215413 | 5 |
| 181 | DA4 | 0.409441611 | 0.417331046 | -0.26853765 | 3 |
| 182 | RMER | 0.408041979 | 0.152770307 | 1.984671951 | 5 |
| 183 | AVHL | 0.407937705 | 0.282648346 | 0.871751431 | 4 |
| 184 | VA7 | 0.402236119 | 0.446115288 | -0.576685065 | 6 |
| 185 | RMEL | 0.394885938 | 0.197530864 | 1.488784708 | 5 |
| 186 | VD2 | 0.394026186 | 0.191012366 | 1.537235413 | 3 |
| 187 | SIADR | 0.387009331 | 0.447222222 | -0.716535893 | 5 |
| 188 | IL1L | 0.385153956 | 0.296052632 | 0.561906148 | 5 |
| 189 | CEPVL | 0.38130192 | 0.392487047 | -0.296755665 | 5 |
| 190 | CEPVR | 0.378950796 | 0.337731959 | 0.151931896 | 5 |
| 191 | IL1R | 0.376533816 | 0.277777778 | 0.644570699 | 5 |
| 192 | URAVR | 0.375411122 | 0 | -0.20098758 | 5 |
| 193 | IL2VR | 0.374876677 | 0 | -0.20098758 | 5 |
| 194 | IL2R | 0.374634623 | 0.444444444 | -0.798705689 | 5 |
| 195 | SIADL | 0.37440658 | 0.322147651 | 0.246458176 | 5 |
| 196 | ASER | 0.369885751 | 0.277042426 | 0.593945505 | 1 |
| 197 | VC3 | 0.369613715 | 0.469886364 | -1.059531217 | 3 |
| 198 | OLQVR | 0.366518775 | 0.32 | 0.197310452 | 5 |
| 199 | RIVL | 0.361920141 | 0.37962963 | -0.352617847 | 5 |
| 200 | AWAR | 0.357515432 | 0.375775434 | -0.357331395 | 1 |
| 201 | SIAVL | 0.357490922 | 0.338823529 | -0.041155645 | 5 |
| 202 | AIYR | 0.354852994 | 0.248457879 | 0.709977171 | 1 |
| 203 | ASJL | 0.353740557 | 0.5 | -1.453274378 | 1 |
| 204 | OLQDL | 0.353269029 | 0.32 | 0.083864907 | 5 |
| 205 | OLQDR | 0.35298403 | 0 | -0.20098758 | 5 |
| 206 | VA9 | 0.351475232 | 0.444444444 | -0.996998529 | 6 |
| 207 | DB2 | 0.347664165 | 0.500825991 | -1.512373214 | 3 |
| 208 | ASGL | 0.337357905 | 0.444818556 | -1.121075554 | 1 |
| 209 | VB11 | 0.336909189 | 0.445919371 | -1.134342777 | 6 |
| 210 | PVM | 0.330227981 | 0.43772482 | -1.121385403 | 4 |
| 211 | VA1 | 0.323157532 | 0.492021277 | -1.646814484 | 3 |
| 212 | DD5 | 0.32032843 | 0.279584639 | 0.147864504 | 6 |
| 213 | VD10 | 0.310794679 | 0.286743269 | 0.004942807 | 6 |
| 214 | AWCR | 0.304949476 | 0.245633388 | 0.306882218 | 1 |
| 215 | VD9 | 0.297524929 | 0.259381607 | 0.125599051 | 6 |
| 216 | ASKR | 0.29665875 | 0.444444444 | -1.466342284 | 1 |

| | | | | | |
|---|---|---|---|---|---|
| 217 | ASEL | 0.282261127 | 0.361111111 | -0.876108391 | 1 |
| 218 | ADFL | 0.279710673 | 0.444444444 | -1.611453277 | 1 |
| 219 | VD8 | 0.266062576 | 0.282418672 | -0.341029974 | 6 |
| 220 | SIBDR | 0.264173524 | 0.5 | -1.220155553 | 5 |
| 221 | URYVR | 0.255295101 | 0.5 | -1.296173428 | 5 |
| 222 | URBL | 0.251369812 | 0.5 | -1.329782113 | 5 |
| 223 | AWCL | 0.249379649 | 0 | -0.20098758 | 1 |
| 224 | URYDL | 0.248174522 | 0.5 | -2.357140475 | 5 |
| 225 | URAVL | 0.241037027 | 0 | -0.20098758 | 5 |
| 226 | URYDR | 0.237559785 | 0 | -0.20098758 | 5 |
| 227 | URBR | 0.234631582 | 0 | -0.20098758 | 5 |
| 228 | VC4 | 0.219372325 | 0.5 | -1.603747531 | 2 |
| 229 | AINR | 0.207559545 | 0.5 | -1.704889644 | 1 |
| 230 | VB5 | 0.201323271 | 0.5 | -1.758285199 | 4 |
| 231 | VB7 | 0.20059122 | 0.5 | -1.764553086 | 4 |
| 232 | AFDR | 0.20001531 | 0.444444444 | -1.293812303 | 1 |
| 233 | VB3 | 0.198212055 | 0.5 | -1.784923716 | 4 |
| 234 | VB6 | 0.197846029 | 0.5 | -1.788057659 | 4 |
| 235 | AFDL | 0.197539343 | 0.444444444 | -1.315011758 | 1 |
| 236 | VB8 | 0.197480004 | 0.5 | -1.791191603 | 6 |
| 237 | VB4 | 0.195466864 | 0.5 | -1.80842829 | 4 |
| 238 | ASJR | 0.184880836 | 0.5 | -1.899066839 | 3 |
| 239 | PHAL | 0.183571038 | 0.5 | -1.910281443 | 4 |
| 240 | PHAR | 0.177172771 | 0.5 | -1.965063997 | 4 |
| 241 | VB2 | 0.158864324 | 0.5 | -1.121822606 | 3 |
| 242 | AIML | 0.140093475 | 0.5 | -1.282540341 | 1 |
| 243 | AIMR | 0 | 0 | -0.20098758 | 1 |
| 244 | AINL | 0 | 0 | -0.20098758 | 1 |
| 245 | ALA | 0 | 0 | -0.20098758 | 3 |
| 246 | ALMR | 0 | 0 | -0.20098758 | 5 |
| 247 | AS7 | 0 | 0 | -0.20098758 | 6 |
| 248 | AS8 | 0 | 0 | -0.20098758 | 6 |
| 249 | AS9 | 0 | 0 | -0.20098758 | 6 |
| 250 | ASIL | 0 | 0 | -0.20098758 | 1 |
| 251 | ASIR | 0 | 0 | -0.20098758 | 1 |
| 252 | AWAL | 0 | 0 | -0.20098758 | 1 |
| 253 | DB1 | 0 | 0 | -0.20098758 | 3 |
| 254 | DB6 | 0 | 0 | -0.20098758 | 6 |
| 255 | IL2DL | 0 | 0 | -0.20098758 | 5 |
| 256 | IL2DR | 0 | 0 | -0.20098758 | 5 |
| 257 | IL2L | 0 | 0 | -0.20098758 | 5 |
| 258 | IL2VL | 0 | 0 | -0.20098758 | 5 |
| 259 | PDA | 0 | 0 | -0.20098758 | 6 |
| 260 | PDB | 0 | 0 | -0.20098758 | 6 |
| 261 | PHCL | 0 | 0 | -0.20098758 | 4 |
| 262 | PHCR | 0 | 0 | -0.20098758 | 4 |
| 263 | PLML | 0 | 0 | -0.20098758 | 3 |
| 264 | PLMR | 0 | 0 | -0.20098758 | 4 |
| 265 | PLNL | 0 | 0 | -0.20098758 | 1 |
| 266 | PLNR | 0 | 0 | -0.20098758 | 1 |
| 267 | PVDL | 0 | 0 | -0.20098758 | 4 |
| 268 | PVDR | 0 | 0 | -0.20098758 | 4 |
| 269 | SDQL | 0 | 0 | -0.20098758 | 5 |
| 270 | SDQR | 0 | 0 | -0.20098758 | 4 |
| 271 | SIBDL | 0 | 0 | -0.20098758 | 5 |
| 272 | URYVL | 0 | 0 | -0.20098758 | 5 |
| 273 | VA8 | 0 | 0 | -0.20098758 | 6 |
| 274 | VB10 | 0 | 0 | -0.20098758 | 6 |
| 275 | VB9 | 0 | 0 | -0.20098758 | 6 |
| 276 | VC1 | 0 | 0 | -0.20098758 | 3 |
| 277 | VC6 | 0 | 0 | -0.20098758 | 6 |

***Supplementary table 3:*** **airport network:** Full set of nodal results for $PC_{norm}$ and PC, and z-score differences between $PC_{norm}$ and PC. Note we used a z-score since $PC_{norm}$ tends to yield higher values than PC -- see Fig. 2 and 3). The last column displays the module community affiliation for each node. See https://en.wikipedia.org/wiki/List_of_airports_by_IATA_code:_A for full naming of specific airports.

| | Node | $PC_{norm}$ | PC | Z-score difference | Module |
|---|---|---|---|---|---|
| 1 | JFK | 0.853633188 | 0.60689782 | -0.400287705 | 3 |
| 2 | PUJ | 0.830163416 | 0.57885355 | -0.371498864 | 3 |
| 3 | YYZ | 0.813662563 | 0.546042274 | -0.268851938 | 3 |
| 4 | YUL | 0.806824245 | 0.49104939 | 0.034200768 | 3 |
| 5 | FRA | 0.781019975 | 0.620267069 | -0.941403954 | 4 |
| 6 | EWR | 0.763787078 | 0.489687189 | -0.228073664 | 3 |
| 7 | YVR | 0.759960001 | 0.461996037 | -0.077889097 | 3 |
| 8 | LHR | 0.740896613 | 0.565572015 | -0.849699454 | 4 |
| 9 | HAV | 0.740439574 | 0.66026847 | -1.448532061 | 1 |
| 10 | LAX | 0.737213446 | 0.48138189 | -0.343042365 | 3 |
| 11 | ANU | 0.73664054 | 0.489695 | -0.398965018 | 3 |
| 12 | AMS | 0.733367541 | 0.575281867 | -0.958189734 | 4 |
| 13 | ORD | 0.73087708 | 0.457365182 | -0.231774091 | 3 |
| 14 | ATL | 0.730516484 | 0.470233554 | -0.31502839 | 3 |
| 15 | AUA | 0.728819252 | 0.497691327 | -0.498510531 | 3 |
| 16 | ANC | 0.724650218 | 0.425703889 | -0.071706744 | 3 |
| 17 | CUN | 0.723621187 | 0.406816339 | 0.040682859 | 3 |
| 18 | CDG | 0.723058694 | 0.5743955 | -1.017488536 | 4 |
| 19 | CCS | 0.721008886 | 0.657781664 | -1.555165546 | 1 |
| 20 | MBJ | 0.706552477 | 0.347093333 | 0.309120538 | 3 |
| 21 | SFO | 0.706492498 | 0.413808219 | -0.111115914 | 3 |
| 22 | HNL | 0.703866349 | 0.4584 | -0.408274061 | 3 |
| 23 | SDQ | 0.703012365 | 0.574502058 | -1.144317362 | 1 |
| 24 | BOS | 0.701639925 | 0.381394276 | 0.062336961 | 3 |
| 25 | BGI | 0.700275671 | 0.450151229 | -0.378959132 | 3 |
| 26 | IAD | 0.699237515 | 0.409190336 | -0.127712061 | 3 |
| 27 | YYT | 0.697515204 | 0.32 | 0.422753337 | 3 |
| 28 | SJO | 0.690855782 | 0.557470679 | -1.113638648 | 1 |
| 29 | MIA | 0.69019384 | 0.475650722 | -0.602884249 | 3 |
| 30 | YHZ | 0.686946028 | 0.301932183 | 0.469944779 | 3 |
| 31 | MEX | 0.683700028 | 0.428855443 | -0.349253702 | 3 |
| 32 | LUX | 0.68362019 | 0.540977366 | -1.055376731 | 4 |
| 33 | SNN | 0.680027083 | 0.408163265 | -0.242146007 | 4 |
| 34 | SXM | 0.67787131 | 0.403077289 | -0.223705263 | 3 |
| 35 | YYC | 0.677839803 | 0.29579 | 0.451291074 | 3 |
| 36 | FCO | 0.674138605 | 0.491871802 | -0.806009846 | 4 |
| 37 | LGW | 0.673991543 | 0.435184807 | -0.450185223 | 4 |
| 38 | KIN | 0.673323422 | 0.38738 | -0.153538367 | 3 |
| 39 | MAN | 0.671852108 | 0.45499811 | -0.588341113 | 4 |
| 40 | PHL | 0.671366346 | 0.328922902 | 0.202035072 | 3 |
| 41 | DKR | 0.670016374 | 0.57897503 | -1.380122016 | 4 |
| 42 | PPT | 0.669404567 | 0.666716049 | -1.936155764 | 2 |
| 43 | GUA | 0.668459495 | 0.397475995 | -0.247686137 | 3 |
| 44 | NRT | 0.665688326 | 0.579870762 | -1.412997007 | 2 |
| 45 | MXP | 0.663255996 | 0.501962111 | -0.937999395 | 4 |
| 46 | LIM | 0.659860919 | 0.615192508 | -1.671962319 | 1 |
| 47 | BOG | 0.658190777 | 0.62934375 | -1.771531557 | 1 |
| 48 | SJU | 0.655545474 | 0.39881893 | -0.337409905 | 3 |
| 49 | AMM | 0.654699496 | 0.543360947 | -1.252384951 | 2 |
| 50 | PTP | 0.653054231 | 0.615306122 | -1.715514087 | 1 |
| 51 | MUC | 0.652570314 | 0.443796433 | -0.639191975 | 4 |
| 52 | SAL | 0.650267437 | 0.379867036 | -0.251355775 | 3 |
| 53 | LOS | 0.650004419 | 0.53888 | -1.253732545 | 2 |
| 54 | TPE | 0.647088557 | 0.569208045 | -1.462947514 | 2 |
| 55 | MAD | 0.645046125 | 0.477911428 | -0.901241222 | 4 |
| 56 | ZRH | 0.644952819 | 0.448945024 | -0.719533207 | 4 |
| 57 | EZE | 0.643612769 | 0.680514178 | -2.185308348 | 1 |
| 58 | ATH | 0.637529017 | 0.434879027 | -0.677731661 | 4 |
| 59 | IAH | 0.637485531 | 0.322421904 | 0.029724769 | 3 |
| 60 | SCL | 0.636552754 | 0.666834977 | -2.143651612 | 1 |
| 61 | TLV | 0.635947426 | 0.435840666 | -0.693737053 | 4 |
| 62 | GDL | 0.634109853 | 0.288597463 | 0.22134898 | 3 |
| 63 | SVO | 0.633321295 | 0.457599683 | -0.847200904 | 4 |
| 64 | ACC | 0.632855216 | 0.495907407 | -1.09121736 | 2 |
| 65 | YOW | 0.630939249 | 0.20244898 | 0.74355639 | 3 |
| 66 | DME | 0.630731409 | 0.505613169 | -1.165664772 | 4 |

| | | | | | |
|---|---|---|---|---|---|
| 67 | JNB | 0.629935391 | 0.565163422 | -1.545443941 | 2 |
| 68 | DXB | 0.627265397 | 0.53632822 | -1.380777578 | 2 |
| 69 | YEG | 0.627125566 | 0.205459566 | 0.700608993 | 3 |
| 70 | SEA | 0.626845434 | 0.245762446 | 0.445206586 | 3 |
| 71 | GLA | 0.624229958 | 0.351869684 | -0.239021643 | 4 |
| 72 | CAI | 0.619015163 | 0.506659408 | -1.245983338 | 2 |
| 73 | DTW | 0.616467299 | 0.246489708 | 0.375316628 | 3 |
| 74 | MCO | 0.61546011 | 0.22948045 | 0.476022972 | 3 |
| 75 | CLO | 0.614503311 | 0.548260355 | -1.536186529 | 1 |
| 76 | POS | 0.614384875 | 0.288262976 | 0.09931816 | 3 |
| 77 | GYE | 0.613677975 | 0.605783934 | -1.90339567 | 1 |
| 78 | GRU | 0.610446384 | 0.654003693 | -2.22719615 | 1 |
| 79 | IST | 0.60923217 | 0.44302663 | -0.907088719 | 4 |
| 80 | BRU | 0.608008059 | 0.380012432 | -0.518223128 | 4 |
| 81 | DFW | 0.607207222 | 0.25516954 | 0.262414804 | 3 |
| 82 | ICN | 0.607152358 | 0.528077916 | -1.455433714 | 2 |
| 83 | DEL | 0.604527279 | 0.530731864 | -1.488656392 | 2 |
| 84 | MGA | 0.602100534 | 0.366826531 | -0.472417886 | 3 |
| 85 | BCN | 0.601929679 | 0.399775047 | -0.680849114 | 4 |
| 86 | ISB | 0.599900709 | 0.544880658 | -1.60681601 | 2 |
| 87 | PEK | 0.59788337 | 0.524086723 | -1.488648642 | 2 |
| 88 | THR | 0.595746981 | 0.509918835 | -1.412930407 | 4 |
| 89 | PTY | 0.594401858 | 0.53565023 | -1.583331954 | 1 |
| 90 | NAS | 0.593763144 | 0.161346483 | 0.768266483 | 3 |
| 91 | FAI | 0.590792882 | 0.132653061 | 0.930150882 | 3 |
| 92 | LAS | 0.589382777 | 0.176970223 | 0.642373972 | 3 |
| 93 | BEY | 0.586119668 | 0.465341784 | -1.192980074 | 2 |
| 94 | PDX | 0.58584412 | 0.159745775 | 0.728503211 | 3 |
| 95 | DAM | 0.585626898 | 0.442614379 | -1.053050115 | 4 |
| 96 | CVG | 0.584808868 | 0.160763393 | 0.715583818 | 3 |
| 97 | UIO | 0.581493537 | 0.604785124 | -2.099657214 | 1 |
| 98 | CPT | 0.580276743 | 0.521137778 | -1.580894307 | 2 |
| 99 | SAN | 0.579629063 | 0.135401916 | 0.842593801 | 3 |
| 100 | DEN | 0.579431318 | 0.158975026 | 0.692995895 | 3 |
| 101 | CLT | 0.579246311 | 0.153949246 | 0.723460495 | 3 |
| 102 | JED | 0.579026866 | 0.503803571 | -1.479670268 | 2 |
| 103 | DUS | 0.57684007 | 0.303305136 | -0.231629109 | 4 |
| 104 | VIE | 0.573485126 | 0.331752041 | -0.431768735 | 4 |
| 105 | PVR | 0.572438829 | 0.099774376 | 1.021559228 | 3 |
| 106 | KOA | 0.572192104 | 0.124444444 | 0.864749556 | 3 |
| 107 | KBP | 0.5707903 | 0.344165183 | -0.526848211 | 4 |
| 108 | LIS | 0.570129046 | 0.357509861 | -0.6149922 | 4 |
| 109 | FLL | 0.57004278 | 0.154775034 | 0.660342646 | 3 |
| 110 | YWG | 0.569946024 | 0.097279429 | 1.021572708 | 3 |
| 111 | CMN | 0.569071254 | 0.355283731 | -0.607639459 | 4 |
| 112 | TIJ | 0.568181222 | 0.129695502 | 0.80646109 | 3 |
| 113 | LHE | 0.567905173 | 0.510005102 | -1.588691082 | 2 |
| 114 | VER | 0.566727442 | 0.177520661 | 0.49633232 | 3 |
| 115 | CPH | 0.565018254 | 0.316813795 | -0.391042222 | 4 |
| 116 | ARN | 0.564741213 | 0.320699085 | -0.417237164 | 4 |
| 117 | MRU | 0.563798069 | 0.4847392 | -1.455531724 | 2 |
| 118 | TPA | 0.563092565 | 0.115225318 | 0.865502161 | 3 |
| 119 | DUB | 0.562989348 | 0.274289632 | -0.136192095 | 4 |
| 120 | DOH | 0.56267188 | 0.479936728 | -1.432395652 | 2 |
| 121 | MLE | 0.562406524 | 0.492561983 | -1.513520561 | 2 |
| 122 | HEL | 0.561578683 | 0.331910431 | -0.507696746 | 4 |
| 123 | BWI | 0.561403831 | 0.127818559 | 0.775620938 | 3 |
| 124 | AUH | 0.561318452 | 0.483810403 | -1.465291553 | 2 |
| 125 | HKG | 0.561306478 | 0.470259695 | -1.380087792 | 2 |
| 126 | LEJ | 0.557918384 | 0.333596735 | -0.541344698 | 4 |
| 127 | ADD | 0.557871375 | 0.472101563 | -1.413297525 | 2 |
| 128 | MTY | 0.557768712 | 0.073525952 | 1.094425372 | 3 |
| 129 | NBO | 0.557692106 | 0.456320499 | -1.315110241 | 2 |
| 130 | BOM | 0.556004751 | 0.476612903 | -1.453436176 | 2 |
| 131 | BKK | 0.555685644 | 0.470869168 | -1.419297185 | 2 |
| 132 | PVG | 0.554187617 | 0.471761342 | -1.434339516 | 2 |
| 133 | ALG | 0.553341239 | 0.331348026 | -0.555998318 | 4 |
| 134 | KEF | 0.552298546 | 0.244897959 | -0.018501291 | 4 |
| 135 | RSW | 0.550612121 | 0.063657943 | 1.111489224 | 3 |
| 136 | PHX | 0.550595074 | 0.089644968 | 0.947836946 | 3 |
| 137 | KHI | 0.54966308 | 0.483908173 | -1.539257985 | 2 |
| 138 | SIN | 0.549356082 | 0.457539571 | -1.375243635 | 2 |
| 139 | FDF | 0.548542024 | 0.571277778 | -2.096159168 | 1 |
| 140 | SLC | 0.547871642 | 0.076394547 | 1.01408679 | 3 |
| 141 | STT | 0.545805263 | 0.069695291 | 1.043243031 | 3 |

| | | | | | |
|---|---|---|---|---|---|
| 142 | MSP | 0.545571496 | 0.090920911 | 0.908191959 | 3 |
| 143 | KWI | 0.544577436 | 0.467057556 | -1.465217094 | 2 |
| 144 | SYR | 0.544498888 | 0.06795728 | 1.045959456 | 3 |
| 145 | LGA | 0.544234184 | 0.091413277 | 0.89667719 | 3 |
| 146 | RDU | 0.543636149 | 0.063954086 | 1.065723382 | 3 |
| 147 | MSY | 0.541812766 | 0.053409578 | 1.12060833 | 3 |
| 148 | TIP | 0.540198455 | 0.332051111 | -0.643134983 | 4 |
| 149 | DCA | 0.539801054 | 0.072266446 | 0.989275406 | 3 |
| 150 | ORY | 0.539631824 | 0.282372191 | -0.334055002 | 4 |
| 151 | ABJ | 0.538950592 | 0.362903047 | -0.845149704 | 2 |
| 152 | MEL | 0.535850583 | 0.460123269 | -1.476498307 | 2 |
| 153 | SAT | 0.534638146 | 0.047551856 | 1.112320646 | 3 |
| 154 | BDL | 0.534598133 | 0.04648038 | 1.11881199 | 3 |
| 155 | MID | 0.534593696 | 0.045207101 | 1.126797237 | 3 |
| 156 | OPO | 0.533875592 | 0.272040763 | -0.30526177 | 4 |
| 157 | MZT | 0.532097059 | 0.017822314 | 1.28342665 | 3 |
| 158 | RUH | 0.532061412 | 0.453827977 | -1.460726456 | 2 |
| 159 | DLA | 0.53191804 | 0.391916667 | -1.07200026 | 2 |
| 160 | BJX | 0.53166471 | 0.018643599 | 1.275537116 | 3 |
| 161 | BIL | 0.531182868 | 0.0129792 | 1.30815267 | 3 |
| 162 | RIC | 0.531143659 | 0.033030965 | 1.181713485 | 3 |
| 163 | ACA | 0.530967142 | 0.017826531 | 1.276289165 | 3 |
| 164 | TLH | 0.530003753 | 0.009648789 | 1.321691493 | 3 |
| 165 | PNS | 0.529612231 | 0.010232422 | 1.315554518 | 3 |
| 166 | BNA | 0.529037237 | 0.045286547 | 1.091328608 | 3 |
| 167 | XNA | 0.52895103 | 0.011646457 | 1.302494362 | 3 |
| 168 | MFE | 0.528869898 | 0.006654064 | 1.333402565 | 3 |
| 169 | SAV | 0.528615395 | 0.010087302 | 1.310194385 | 3 |
| 170 | HSV | 0.528504571 | 0.009443787 | 1.313546783 | 3 |
| 171 | FAT | 0.528393986 | 0.001814059 | 1.360867252 | 3 |
| 172 | EMA | 0.528293399 | 0.228688264 | -0.067560663 | 4 |
| 173 | TYS | 0.528279964 | 0.008994646 | 1.314959854 | 3 |
| 174 | SMF | 0.528093737 | 0.025246079 | 1.211512189 | 3 |
| 175 | MEM | 0.528007976 | 0.029318534 | 1.185343148 | 3 |
| 176 | LEX | 0.527979087 | 0.001141869 | 1.362486477 | 3 |
| 177 | BZN | 0.527878508 | 0 | 1.369039658 | 3 |
| 178 | CLE | 0.527785989 | 0.036158416 | 1.140900456 | 3 |
| 179 | CUU | 0.527747744 | 0.022663265 | 1.225589243 | 3 |
| 180 | SJD | 0.527739206 | 0.007603175 | 1.320313674 | 3 |
| 181 | FSD | 0.527705384 | 0 | 1.367950127 | 3 |
| 182 | CID | 0.527691089 | 0 | 1.367860165 | 3 |
| 183 | BTR | 0.527672009 | 0 | 1.367740088 | 3 |
| 184 | FNT | 0.527599129 | 0 | 1.367281433 | 3 |
| 185 | CRP | 0.52758151 | 0 | 1.367170547 | 3 |
| 186 | PVD | 0.527506647 | 0.030069197 | 1.177463945 | 3 |
| 187 | IND | 0.527360659 | 0.026371165 | 1.199818138 | 3 |
| 188 | MLI | 0.527338989 | 0 | 1.365644283 | 3 |
| 189 | ICT | 0.527274871 | 0.004099723 | 1.339439846 | 3 |
| 190 | OGG | 0.527271604 | 0 | 1.365220209 | 3 |
| 191 | MYR | 0.527261335 | 0 | 1.365155581 | 3 |
| 192 | FAR | 0.527208016 | 0 | 1.36482003 | 3 |
| 193 | VPS | 0.527176809 | 0.001426612 | 1.35564549 | 3 |
| 194 | LGB | 0.527108723 | 0 | 1.364195143 | 3 |
| 195 | SDF | 0.527075829 | 0.028288281 | 1.185960564 | 3 |
| 196 | OMA | 0.526921951 | 0.026824229 | 1.194205922 | 3 |
| 197 | AKL | 0.5268456 | 0.495513672 | -1.755893248 | 2 |
| 198 | HPN | 0.526783256 | 0 | 1.362146869 | 3 |
| 199 | GRB | 0.526736534 | 0 | 1.361852834 | 3 |
| 200 | MDT | 0.526701203 | 0 | 1.361630485 | 3 |
| 201 | PSP | 0.526654239 | 0 | 1.361334924 | 3 |
| 202 | SBA | 0.52661251 | 0 | 1.361072309 | 3 |
| 203 | GSP | 0.526563387 | 0.007871435 | 1.311225603 | 3 |
| 204 | COS | 0.526532179 | 0.000975 | 1.35443076 | 3 |
| 205 | BTV | 0.526394507 | 0 | 1.359700347 | 3 |
| 206 | DSM | 0.526161817 | 0 | 1.358235948 | 3 |
| 207 | GSO | 0.526067814 | 0.006894965 | 1.314252045 | 3 |
| 208 | MSN | 0.525952277 | 0 | 1.356917245 | 3 |
| 209 | SBN | 0.525939447 | 0 | 1.356836501 | 3 |
| 210 | SRQ | 0.525930743 | 0 | 1.356781726 | 3 |
| 211 | CAE | 0.525887109 | 0 | 1.35650712 | 3 |
| 212 | CHS | 0.525833056 | 0.006143347 | 1.317504818 | 3 |
| 213 | BUF | 0.525792577 | 0.025655845 | 1.194451426 | 3 |
| 214 | YYJ | 0.525718444 | 0 | 1.355445652 | 3 |
| 215 | PWM | 0.525377804 | 0 | 1.353301893 | 3 |
| 216 | ISP | 0.525209878 | 0 | 1.352245082 | 3 |

| | | | | | |
|---|---|---|---|---|---|
| 217 | GRR | 0.525007404 | 0 | 1.350970845 | 3 |
| 218 | DAY | 0.52481538 | 0.003742604 | 1.32620892 | 3 |
| 219 | EUG | 0.524808744 | 0 | 1.349720608 | 3 |
| 220 | HMO | 0.524743471 | 0.020153061 | 1.222479904 | 3 |
| 221 | MAF | 0.524589243 | 0 | 1.348339216 | 3 |
| 222 | LBB | 0.524489163 | 0 | 1.347709378 | 3 |
| 223 | AMA | 0.524440283 | 0 | 1.347401765 | 3 |
| 224 | HRL | 0.524245586 | 0 | 1.346176467 | 3 |
| 225 | ROC | 0.524087751 | 0 | 1.345183158 | 3 |
| 226 | BOI | 0.523690802 | 0.001206525 | 1.335091965 | 3 |
| 227 | AUS | 0.523592064 | 0.024067924 | 1.190596185 | 3 |
| 228 | TUS | 0.523471729 | 0.008158861 | 1.289959899 | 3 |
| 229 | STL | 0.523057915 | 0.029248438 | 1.154631908 | 3 |
| 230 | BUR | 0.52302258 | 0.000677765 | 1.334214289 | 3 |
| 231 | GEG | 0.52290546 | 0 | 1.337742609 | 3 |
| 232 | CJS | 0.522874675 | 0.020958678 | 1.205648933 | 3 |
| 233 | LIH | 0.522819092 | 0 | 1.337199066 | 3 |
| 234 | SNA | 0.522811663 | 0.010168006 | 1.273161672 | 3 |
| 235 | PBI | 0.522808102 | 0 | 1.337129905 | 3 |
| 236 | ALB | 0.522699816 | 0.00519873 | 1.303731081 | 3 |
| 237 | TUN | 0.522399999 | 0.275165406 | -0.39714592 | 4 |
| 238 | JAN | 0.522035545 | 0.000644988 | 1.328208821 | 3 |
| 239 | RNO | 0.521971175 | 0.000634755 | 1.327868116 | 3 |
| 240 | LIT | 0.521917055 | 0.000624843 | 1.327589905 | 3 |
| 241 | DAL | 0.520640193 | 0.000596878 | 1.319730181 | 3 |
| 242 | MKE | 0.520589177 | 0.002687183 | 1.306254129 | 3 |
| 243 | ONT | 0.52038151 | 0.003197598 | 1.301735002 | 3 |
| 244 | ELP | 0.520142886 | 0 | 1.320356812 | 3 |
| 245 | CUL | 0.520108617 | 0.0249375 | 1.163201157 | 3 |
| 246 | YLW | 0.520091472 | 0 | 1.32003325 | 3 |
| 247 | MHT | 0.519917078 | 0.004332602 | 1.291669216 | 3 |
| 248 | MDW | 0.519913152 | 0.00989282 | 1.256652212 | 3 |
| 249 | BHM | 0.519878602 | 0.004224323 | 1.292108517 | 3 |
| 250 | HOU | 0.519861263 | 0 | 1.318584467 | 3 |
| 251 | JAX | 0.519855869 | 0.004845336 | 1.288057203 | 3 |
| 252 | VSA | 0.519778702 | 0.0219 | 1.180240891 | 3 |
| 253 | PIT | 0.519657577 | 0.017625573 | 1.206379009 | 3 |
| 254 | ORF | 0.519308417 | 0.004784024 | 1.284997765 | 3 |
| 255 | TUL | 0.519219663 | 0 | 1.314546661 | 3 |
| 256 | OKC | 0.519046363 | 0.002024984 | 1.300712131 | 3 |
| 257 | SJC | 0.518082596 | 0.000963268 | 1.301328558 | 3 |
| 258 | ABQ | 0.517263843 | 0.000470522 | 1.299276886 | 3 |
| 259 | OAK | 0.517030269 | 0.000919031 | 1.294984308 | 3 |
| 260 | MCI | 0.515543005 | 0.007817762 | 1.242208447 | 3 |
| 261 | STN | 0.515413105 | 0.21736263 | -0.077344655 | 4 |
| 262 | CMH | 0.515314534 | 0.003448613 | 1.268267117 | 3 |
| 263 | PRG | 0.512646511 | 0.197140788 | 0.032507036 | 4 |
| 264 | KRT | 0.511562432 | 0.433185596 | -1.459823983 | 2 |
| 265 | JNU | 0.511365239 | 0 | 1.265116155 | 3 |
| 266 | EDI | 0.511097491 | 0.176855056 | 0.150423396 | 4 |
| 267 | SAH | 0.510726485 | 0.434321361 | -1.472232628 | 2 |
| 268 | OTP | 0.509987733 | 0.208818182 | -0.057715272 | 4 |
| 269 | MDE | 0.507787248 | 0.476946746 | -1.758985956 | 1 |
| 270 | GVA | 0.507028551 | 0.209686166 | -0.081800901 | 4 |
| 271 | LCA | 0.505756365 | 0.23154479 | -0.227370792 | 4 |
| 272 | SYD | 0.505702161 | 0.432394464 | -1.491725761 | 2 |
| 273 | TXL | 0.505156259 | 0.194461813 | 0.002228064 | 4 |
| 274 | BFS | 0.504929006 | 0.165289256 | 0.184390494 | 4 |
| 275 | DAR | 0.504748372 | 0.427555556 | -1.467275414 | 2 |
| 276 | SHJ | 0.50397869 | 0.415764938 | -1.397917001 | 2 |
| 277 | VCE | 0.503131107 | 0.168628166 | 0.152062846 | 4 |
| 278 | ITO | 0.503036956 | 0 | 1.212703502 | 3 |
| 279 | CGN | 0.495071791 | 0.180208148 | 0.028466207 | 4 |
| 280 | BUD | 0.494885017 | 0.178245825 | 0.03964033 | 4 |
| 281 | MRS | 0.494304634 | 0.16861678 | 0.096586568 | 4 |
| 282 | KTN | 0.493564418 | 0 | 1.153089668 | 3 |
| 283 | GIG | 0.492964201 | 0.530649796 | -2.190243493 | 1 |
| 284 | NTE | 0.490282283 | 0.196270408 | -0.102760907 | 4 |
| 285 | LED | 0.489744926 | 0.193761815 | -0.09035526 | 4 |
| 286 | GOT | 0.487683558 | 0.171088156 | 0.039364748 | 4 |
| 287 | KIX | 0.483279843 | 0.378138184 | -1.291384042 | 2 |
| 288 | NAN | 0.482907438 | 0.397338843 | -1.414563853 | 2 |
| 289 | NGO | 0.481496695 | 0.379060764 | -1.30841209 | 2 |
| 290 | MAA | 0.481193092 | 0.3871025 | -1.360932085 | 2 |
| 291 | BHX | 0.480177269 | 0.141078343 | 0.180986909 | 4 |

| | | | | | |
|---|---|---|---|---|---|
| 292 | KRK | 0.479678092 | 0.121928166 | 0.29836386 | 4 |
| 293 | VVI | 0.477749255 | 0.51752 | -2.203366035 | 1 |
| 294 | CWL | 0.47707454 | 0.120707596 | 0.289660292 | 4 |
| 295 | BAH | 0.475560987 | 0.375428701 | -1.322909693 | 2 |
| 296 | LYS | 0.47355634 | 0.120027778 | 0.271797413 | 4 |
| 297 | BSL | 0.472879701 | 0.148433153 | 0.088774618 | 4 |
| 298 | BLQ | 0.472676645 | 0.123766541 | 0.242731923 | 4 |
| 299 | WAW | 0.471799355 | 0.103584162 | 0.364225268 | 4 |
| 300 | DMM | 0.471430306 | 0.369556787 | -1.311951538 | 2 |
| 301 | MNL | 0.46865091 | 0.35544186 | -1.240613273 | 2 |
| 302 | SUF | 0.468023408 | 0.104938272 | 0.331940118 | 4 |
| 303 | OSL | 0.46787529 | 0.118248521 | 0.247242135 | 4 |
| 304 | PSA | 0.467446935 | 0.097304405 | 0.376354641 | 4 |
| 305 | SGN | 0.465465356 | 0.367220703 | -1.334789197 | 2 |
| 306 | HAM | 0.464652734 | 0.096578433 | 0.36333859 | 4 |
| 307 | NCE | 0.461019932 | 0.092300219 | 0.367400383 | 4 |
| 308 | KUL | 0.459827138 | 0.34894727 | -1.255271582 | 2 |
| 309 | BOD | 0.457223924 | 0.087621224 | 0.372957308 | 4 |
| 310 | NCL | 0.456460982 | 0.087693889 | 0.367698565 | 4 |
| 311 | AMD | 0.453698347 | 0.361259259 | -1.371325555 | 2 |
| 312 | HKT | 0.452542819 | 0.3791875 | -1.491426063 | 2 |
| 313 | LTN | 0.451028649 | 0.074504421 | 0.416516793 | 4 |
| 314 | BEG | 0.450264714 | 0.091402344 | 0.305364843 | 4 |
| 315 | AGP | 0.449079552 | 0.071611601 | 0.422455956 | 4 |
| 316 | CMB | 0.44879611 | 0.338685172 | -1.260110714 | 2 |
| 317 | PMO | 0.4471646 | 0.060546875 | 0.480038532 | 4 |
| 318 | TFN | 0.445529042 | 0.110615917 | 0.154644269 | 4 |
| 319 | TLS | 0.445001925 | 0.055510204 | 0.498125546 | 4 |
| 320 | STR | 0.444255804 | 0.049351066 | 0.532191464 | 4 |
| 321 | LPL | 0.443866272 | 0.052593134 | 0.509336595 | 4 |
| 322 | MVD | 0.443667547 | 0.492663265 | -2.261421869 | 1 |
| 323 | SOF | 0.443531965 | 0.069038399 | 0.403737159 | 4 |
| 324 | SCQ | 0.443447883 | 0.099623269 | 0.21072724 | 4 |
| 325 | RIX | 0.439486782 | 0.042533081 | 0.545086268 | 4 |
| 326 | SXF | 0.438010101 | 0.048895508 | 0.495752154 | 4 |
| 327 | FAO | 0.437588137 | 0.038446751 | 0.558854097 | 4 |
| 328 | NAP | 0.436177674 | 0.035702479 | 0.5672482 | 4 |
| 329 | BRS | 0.435270468 | 0.033888228 | 0.572956534 | 4 |
| 330 | FNC | 0.434069604 | 0.058710744 | 0.409182749 | 4 |
| 331 | MLA | 0.430182317 | 0.032295482 | 0.550958792 | 4 |
| 332 | BLR | 0.429720651 | 0.310866941 | -1.205089546 | 2 |
| 333 | ALC | 0.428673673 | 0.030295858 | 0.554048691 | 4 |
| 334 | LPA | 0.422115553 | 0.009532562 | 0.643446594 | 4 |
| 335 | PMI | 0.419817173 | 0 | 0.688973707 | 4 |
| 336 | IBZ | 0.41912439 | 0 | 0.684613791 | 4 |
| 337 | ACE | 0.418771855 | 0 | 0.68239517 | 4 |
| 338 | HAJ | 0.418744975 | 0 | 0.682226006 | 4 |
| 339 | TFS | 0.418601211 | 0 | 0.681321252 | 4 |
| 340 | FUE | 0.418547057 | 0 | 0.680980443 | 4 |
| 341 | LBA | 0.418318465 | 0 | 0.679541836 | 4 |
| 342 | VLC | 0.418225729 | 0 | 0.678958219 | 4 |
| 343 | HER | 0.418214733 | 0 | 0.678889019 | 4 |
| 344 | MAH | 0.418143589 | 0 | 0.678441283 | 4 |
| 345 | BGY | 0.418051982 | 0 | 0.677864774 | 4 |
| 346 | SVQ | 0.41780421 | 0 | 0.676305459 | 4 |
| 347 | NUE | 0.417710432 | 0 | 0.675715283 | 4 |
| 348 | CTA | 0.417679609 | 0 | 0.675521304 | 4 |
| 349 | ORK | 0.417452397 | 0 | 0.674091387 | 4 |
| 350 | GRO | 0.417339674 | 0 | 0.673381982 | 4 |
| 351 | SKG | 0.417229108 | 0 | 0.672686153 | 4 |
| 352 | LCY | 0.417045738 | 0 | 0.671532146 | 4 |
| 353 | BRE | 0.417019633 | 0 | 0.671367854 | 4 |
| 354 | SOU | 0.416581533 | 0 | 0.66861075 | 4 |
| 355 | VNO | 0.416562284 | 0 | 0.668489609 | 4 |
| 356 | ABZ | 0.416484565 | 0 | 0.668000498 | 4 |
| 357 | BLL | 0.416452811 | 0 | 0.667800658 | 4 |
| 358 | TRN | 0.416437843 | 0 | 0.667706461 | 4 |
| 359 | RAK | 0.416420079 | 0 | 0.667594666 | 4 |
| 360 | LIN | 0.416373657 | 0 | 0.667302512 | 4 |
| 361 | AYT | 0.416344988 | 0 | 0.667122092 | 4 |
| 362 | BIO | 0.416165736 | 0 | 0.665994001 | 4 |
| 363 | SZG | 0.415899705 | 0 | 0.664319775 | 4 |
| 364 | SVG | 0.415881738 | 0 | 0.664206704 | 4 |
| 365 | BGO | 0.415852701 | 0 | 0.664023966 | 4 |
| 366 | JER | 0.415764901 | 0 | 0.663471411 | 4 |

| | | | | | |
|---|---|---|---|---|---|
| 367 | VRN | 0.415680446 | 0 | 0.662939909 | 4 |
| 368 | LJU | 0.415645762 | 0 | 0.662721631 | 4 |
| 369 | TLL | 0.415504428 | 0 | 0.661832167 | 4 |
| 370 | ESB | 0.415373354 | 0 | 0.661007276 | 4 |
| 371 | ZAG | 0.415353412 | 0 | 0.660881773 | 4 |
| 372 | FLR | 0.415279616 | 0 | 0.660417351 | 4 |
| 373 | XRY | 0.415245059 | 0 | 0.660199874 | 4 |
| 374 | ADB | 0.414846401 | 0 | 0.657690987 | 4 |
| 375 | DRS | 0.414806941 | 0 | 0.657442649 | 4 |
| 376 | SXB | 0.414523077 | 0 | 0.655656197 | 4 |
| 377 | BRI | 0.413861292 | 0 | 0.651491366 | 4 |
| 378 | GRZ | 0.413386331 | 0 | 0.648502275 | 4 |
| 379 | MCT | 0.41313852 | 0.275624567 | -1.087654424 | 2 |
| 380 | DAC | 0.41303396 | 0.293398866 | -1.200172035 | 2 |
| 381 | BHD | 0.413017868 | 0 | 0.646183419 | 4 |
| 382 | CAN | 0.412417164 | 0.257873136 | -0.980478491 | 2 |
| 383 | GOA | 0.412363018 | 0 | 0.642062227 | 4 |
| 384 | MME | 0.412042443 | 0 | 0.640044743 | 4 |
| 385 | TRD | 0.411411461 | 0 | 0.636073761 | 4 |
| 386 | MPL | 0.410258782 | 0 | 0.62881957 | 4 |
| 387 | OVD | 0.409629862 | 0 | 0.624861568 | 4 |
| 388 | GUM | 0.409577055 | 0.278037037 | -1.125250384 | 2 |
| 389 | SSA | 0.408612188 | 0.426987654 | -2.068718426 | 1 |
| 390 | VGO | 0.407947013 | 0 | 0.614270837 | 4 |
| 391 | HAN | 0.407880056 | 0.297855372 | -1.260653545 | 2 |
| 392 | TRS | 0.407599454 | 0 | 0.612083535 | 4 |
| 393 | CCU | 0.40374626 | 0.281024691 | -1.18074782 | 2 |
| 394 | HYD | 0.403050939 | 0.260839506 | -1.058091618 | 2 |
| 395 | LCG | 0.399505896 | 0 | 0.561148082 | 4 |
| 396 | TOS | 0.396632475 | 0 | 0.543064686 | 4 |
| 397 | PEN | 0.395324211 | 0.280969136 | -1.233400951 | 2 |
| 398 | BOO | 0.394138644 | 0 | 0.527370176 | 4 |
| 399 | SZX | 0.373210484 | 0.197008601 | -0.844178402 | 2 |
| 400 | NKG | 0.36372967 | 0.177492063 | -0.781020253 | 2 |
| 401 | SYX | 0.361187138 | 0.199890261 | -0.937980564 | 2 |
| 402 | XMN | 0.356439359 | 0.158442734 | -0.707016839 | 2 |
| 403 | TSN | 0.354600016 | 0.168962351 | -0.784795883 | 2 |
| 404 | REC | 0.349515888 | 0.349246914 | -1.951382758 | 1 |
| 405 | BNE | 0.347291894 | 0.160208333 | -0.775696383 | 2 |
| 406 | MHD | 0.346263668 | 0.20428125 | -1.059532886 | 2 |
| 407 | CTS | 0.343855871 | 0.152777778 | -0.750557459 | 2 |
| 408 | FOR | 0.341821651 | 0.33465374 | -1.907965448 | 1 |
| 409 | MFM | 0.339709248 | 0.140268861 | -0.697930764 | 2 |
| 410 | PUS | 0.33681054 | 0.144970414 | -0.745761735 | 2 |
| 411 | SHE | 0.335151558 | 0.121321865 | -0.607374072 | 2 |
| 412 | CGK | 0.332858089 | 0.121216049 | -0.621141699 | 2 |
| 413 | GOI | 0.332434237 | 0.146708333 | -0.784240571 | 2 |
| 414 | KMQ | 0.328101353 | 0.18 | -1.021024393 | 2 |
| 415 | URC | 0.327274544 | 0.120025712 | -0.648789616 | 2 |
| 416 | DPS | 0.311506246 | 0.109722222 | -0.683181475 | 2 |
| 417 | NAT | 0.311477042 | 0.29722314 | -1.863370957 | 1 |
| 418 | CWB | 0.302522944 | 0.269135734 | -1.742958674 | 1 |
| 419 | TNA | 0.299811808 | 0.058769513 | -0.436116111 | 2 |
| 420 | CTU | 0.299080699 | 0.04875 | -0.377661087 | 2 |
| 421 | DLC | 0.299063307 | 0.04875 | -0.377770543 | 2 |
| 422 | PER | 0.29013716 | 0.052930748 | -0.460256595 | 2 |
| 423 | KMG | 0.281539496 | 0.001865702 | -0.192995229 | 2 |
| 424 | COK | 0.278033828 | 0.004686981 | -0.232812807 | 2 |
| 425 | TAO | 0.27771705 | 0 | -0.205309669 | 2 |
| 426 | HGH | 0.277348942 | 0 | -0.207626291 | 2 |
| 427 | CKG | 0.277118376 | 0 | -0.209077322 | 2 |
| 428 | TRV | 0.276465346 | 0.005302469 | -0.246557262 | 2 |
| 429 | XIY | 0.276438758 | 0 | -0.213354385 | 2 |
| 430 | WUH | 0.276080816 | 0 | -0.215607029 | 2 |
| 431 | KWL | 0.275623604 | 0 | -0.218484418 | 2 |
| 432 | HAK | 0.275453391 | 0 | -0.219555626 | 2 |
| 433 | FOC | 0.274909452 | 0 | -0.222978816 | 2 |
| 434 | CGO | 0.274813405 | 0 | -0.223583271 | 2 |
| 435 | CSX | 0.274673622 | 0 | -0.224462972 | 2 |
| 436 | FUK | 0.274397759 | 0 | -0.226199068 | 2 |
| 437 | KWE | 0.274190433 | 0 | -0.227503842 | 2 |
| 438 | TYN | 0.273730836 | 0 | -0.230396236 | 2 |
| 439 | HRB | 0.273718277 | 0 | -0.230475275 | 2 |
| 440 | WNZ | 0.272839124 | 0 | -0.236008077 | 2 |
| 441 | LHW | 0.272378391 | 0 | -0.238907622 | 2 |

| | | | | | |
|---|---|---|---|---|---|
| 442 | DUR | 0.272214021 | 0.022125 | -0.379182051 | 2 |
| 443 | NGB | 0.27212233 | 0 | -0.240519101 | 2 |
| 444 | CGQ | 0.271720238 | 0 | -0.243049601 | 2 |
| 445 | SHA | 0.271378811 | 0 | -0.245198317 | 2 |
| 446 | NNG | 0.269944841 | 0 | -0.254222763 | 2 |
| 447 | OKA | 0.268918822 | 0 | -0.260679844 | 2 |
| 448 | YNT | 0.268271702 | 0 | -0.264752388 | 2 |
| 449 | HND | 0.265534199 | 0 | -0.281980401 | 2 |
| 450 | KHH | 0.264611624 | 0 | -0.287786474 | 2 |
| 451 | SDJ | 0.262595336 | 0 | -0.300475649 | 2 |
| 452 | CJU | 0.260387426 | 0 | -0.314370756 | 2 |
| 453 | ADL | 0.26036022 | 0.001183432 | -0.321989703 | 2 |
| 454 | BKI | 0.26023124 | 0 | -0.315353686 | 2 |
| 455 | CNS | 0.26018389 | 0.001467456 | -0.324886863 | 2 |
| 456 | GAU | 0.260033118 | 0.011944444 | -0.391770901 | 2 |
| 457 | PNQ | 0.25911161 | 0.008493827 | -0.375854372 | 2 |
| 458 | RGN | 0.257954748 | 0.01376 | -0.41627667 | 2 |
| 459 | CEB | 0.257832313 | 0.007107438 | -0.375180407 | 2 |
| 460 | HIJ | 0.257636372 | 0 | -0.331684059 | 2 |
| 461 | PNH | 0.257461771 | 0 | -0.332782876 | 2 |
| 462 | ITM | 0.256430149 | 0 | -0.339275221 | 2 |
| 463 | KOJ | 0.251296939 | 0 | -0.37158022 | 2 |
| 464 | CNX | 0.250035427 | 0 | -0.379519335 | 2 |
| 465 | DRW | 0.250005095 | 0.009728395 | -0.440934254 | 2 |
| 466 | CHC | 0.249438717 | 0.010246914 | -0.447761865 | 2 |
| 467 | KCH | 0.245937096 | 0 | -0.405311498 | 2 |
| 468 | KMI | 0.245847869 | 0 | -0.405873028 | 2 |
| 469 | NGS | 0.245805876 | 0 | -0.406137306 | 2 |
| 470 | OOL | 0.245761128 | 0.000277778 | -0.408167068 | 2 |
| 471 | SUB | 0.245195239 | 0 | -0.40998025 | 2 |
| 472 | KMJ | 0.244762453 | 0 | -0.412703915 | 2 |
| 473 | UKB | 0.244662042 | 0 | -0.413335835 | 2 |
| 474 | MYJ | 0.244644245 | 0 | -0.413447838 | 2 |
| 475 | WLG | 0.243958954 | 0.000244898 | -0.419301827 | 2 |
| 476 | CBR | 0.2371014 | 0.00048 | -0.463938273 | 2 |
| 477 | BEL | 0.223963816 | 0.111122449 | -1.242927221 | 1 |
| 478 | MAO | 0.221339668 | 0.099210938 | -1.18447875 | 1 |
| 479 | BSB | 0.220491664 | 0.094905 | -1.162716823 | 1 |
| 480 | PLZ | 0.21769923 | 0.009777778 | -0.644556598 | 2 |
| 481 | HBA | 0.207213939 | 0.000375 | -0.651369214 | 2 |
| 482 | ISG | 0.206673859 | 0 | -0.652408113 | 2 |
| 483 | HKD | 0.205542433 | 0 | -0.659528557 | 2 |
| 484 | MES | 0.203874453 | 0 | -0.670025708 | 2 |
| 485 | CGH | 0.177394761 | 0 | -0.836671222 | 1 |
| 486 | POA | 0.175706276 | 0 | -0.847297425 | 1 |
| 487 | CNF | 0.172988394 | 0 | -0.864401959 | 1 |
| 488 | SLZ | 0.166598674 | 0 | -0.904614593 | 1 |
| 489 | CPQ | 0.166475257 | 0 | -0.905391299 | 1 |
| 490 | FLN | 0.166394732 | 0 | -0.905898071 | 1 |
| 491 | VIX | 0.163755796 | 0 | -0.922505773 | 1 |
| 492 | CGB | 0.161801185 | 0 | -0.93480679 | 1 |
| 493 | GYN | 0.158107259 | 0 | -0.958053895 | 1 |
| 494 | GMP | 0.15246357 | 0 | -0.993571507 | 2 |
| 495 | UPG | 0.151232581 | 0 | -1.001318531 | 2 |
| 496 | MZG | 0.130842854 | 0 | -1.129637872 | 2 |
| 497 | TSA | 0.1304326 | 0 | -1.132219737 | 2 |
| 498 | DMK | 0.130257917 | 0 | -1.133319072 | 2 |
| 499 | SDU | 0.001 | 0 | -1.946782169 | 1 |
